# *LEAFY* maintains apical stem cell activity during shoot development in the fern *Ceratopteris richardii*

**DOI:** 10.1101/360107

**Authors:** Andrew R.G. Plackett, Stephanie J. Conway, Kristen D. Hewett Hazelton, Ester H. Rabbinowitsch, Jane. A. Langdale, Verónica S. Di Stilio

**Author notes:** Department of Plant Sciences, University of Cambridge, Downing Street, Cambridge, CB2 3EA, UK. Department of Organismic and Evolutionary Biology, Harvard University, 16 Divinity Ave., Cambridge, MA 02138, USA. Equal contribution (Andrew R.G. Plackett, Stephanie J. Conway). Equal contribution (Jane. A. Langdale, Verónica S. Di Stilio). **COMPETING INTERESTS** The authors declare that they have no competing interests.

## Abstract

During land plant evolution, determinate spore-bearing axes (retained in extant bryophytes such as mosses) were progressively transformed into indeterminate branching shoots with specialized reproductive axes that form flowers. The LEAFY transcription factor, which is required for the first zygotic cell division in mosses and primarily for floral meristem identity in flowering plants, may have facilitated developmental innovations during these transitions. Mapping the LEAFY evolutionary trajectory has been challenging, however, because there is no functional overlap between mosses and flowering plants, and no functional data from intervening lineages. Here, we report a transgenic analysis in the fern *Ceratopteris richardii* that reveals a role for LEAFY in maintaining cell divisions in the apical stem cells of both haploid and diploid phases of the lifecycle. These results support an evolutionary trajectory in which an ancestral LEAFY module that promotes cell proliferation was progressively co-opted, adapted and specialized as novel shoot developmental contexts emerged.

## INTRODUCTION

Land plants are characterized by the alternation of haploid (gametophyte) and diploid (sporophyte) phases within their lifecycle, both of which are multicellular (Niklas and Kutschera, 2010; Bowman et al., 2016). In the earliest diverging bryophyte lineages (liverworts, mosses and hornworts) the free-living indeterminate gametophyte predominates the lifecycle, producing gametes that fuse to form the sporophyte. The sporophyte embryo develops on the surface of the gametophyte, ultimately forming a simple determinate spore-producing axis (Kato and Akiyama, 2005; Ligrone et al., 2012). By contrast, angiosperm (flowering plant) sporophytes range from small herbaceous to large arborescent forms, all developing from an indeterminate vegetative shoot apex that ultimately transitions to flowering; and gametophytes are few-celled determinate structures produced within flowers (Schmidt et al., 2015). A series of developmental innovations during the course of land plant evolution thus simplified gametophyte form whilst increasing sporophyte complexity, with a prolonged and plastic phase of vegetative development arising in the sporophyte of all vascular plants (lycophytes, ferns, gymnosperms and angiosperms).

Studies aimed at understanding how gene function evolved to facilitate developmental innovations during land plant evolution have thus far largely relied on comparative analyses between bryophytes and angiosperms, lineages that diverged over 450 million years ago. Such comparisons have revealed examples of both sub- and neo-functionalization following gene duplication, and of co-option of existing gene regulatory networks into new developmental contexts. For example, a single bHLH transcription factor in the moss *Physcomitrella patens* regulates stomatal differentiation, whereas gene duplications have resulted in three homologs with sub-divided stomatal patterning roles in the angiosperm *Arabidopsis thaliana* (hereafter ‘Arabidopsis’) (MacAlister and Bergmann, 2011); class III HD-ZIP transcription factors play a conserved role in the regulation of leaf polarity in *P. patens* and Arabidopsis but gene family members have acquired regulatory activity in meristems of angiosperms (Yip et al., 2016); and the gene regulatory network that produces rhizoids on the gametophytes of both the moss *P. patens* and the liverwort *Marchantia polymorpha* has been co-opted to regulate root hair formation in Arabidopsis sporophytes (Menand et al., 2007; Pires et al., 2013; Proust et al., 2016). In many cases, however, interpreting the evolutionary trajectory of gene function by comparing lineages as disparate as bryophytes and angiosperms has proved challenging, particularly when only a single representative gene remains in most extant taxa – as is the case for the *LEAFY* (*LFY*) gene family (Himi et al., 2001; Maizel et al., 2005; Sayou et al., 2014).

The LFY transcription factor, which is present across all extant land plant lineages and related streptophyte algae (Sayou et al., 2014), has distinct functional roles in bryophytes and angiosperms. In *P. patens*, LFY regulates cell divisions during sporophyte development (including the first division of the zygote) (Tanahashi et al., 2005), whereas in angiosperms the major role is to promote the transition from inflorescence to floral meristem identity (Carpenter and Coen, 1990; Schultz and Haughn, 1991; Weigel et al., 1992; Weigel and Nilsson, 1995; Souer et al., 1998; Molinero-Rosales et al., 1999). Given that LFY proteins from liverworts and all vascular plant lineages tested to date (ferns, gymnosperms and angiosperms) bind a conserved target DNA motif, whereas hornwort and moss homologs bind to different lineage-specific motifs (Sayou et al., 2014), the divergent roles in mosses and angiosperms may have arisen through the activation of distinct networks of downstream targets. This suggestion is supported by the observation that PpLFY cannot complement loss-of-function *lfy* mutants in Arabidopsis (Maizel, 2005). Similar complementation studies indicate progressive functional changes as vascular plant lineages diverged in that the *lfy* mutant is not complemented by lycophyte LFY proteins (Yang et al., 2017) but is partially complemented by fern and (increasingly) gymnosperm homologs (Maizel et al., 2005). Because LFY proteins from ferns, gymnosperms and angiosperms recognize the same DNA motif, this progression likely reflects co-option of a similar *LFY* gene regulatory network into different developmental contexts. As such, the role in floral meristem identity in angiosperms would have been co-opted from an unknown ancestral context in non-flowering vascular plants, a context that cannot be predicted from existing bryophyte data.

The role of *LFY* in non-flowering vascular plant lineages has thus far been hypothesized on the basis of expression patterns in the lycophyte *Isoetes sinensis* (Yang et al., 2017), the fern *Ceratopteris richardii* (hereafter ‘Ceratopteris’) (Himi et al., 2001) and several gymnosperm species (Mellerowicz et al.; Mouradov et al., 1998; Shindo et al., 2001; Carlsbecker et al., 2004; Vázquez-Lobo et al., 2007; Carlsbecker et al., 2013). These studies reported broad expression in vegetative and reproductive sporophyte tissues of *I. sinensis* and gymnosperms, and in both gametophytes and sporophytes of Ceratopteris. Although gene expression can be indicative of potential roles in each case, the possible evolutionary trajectories and differing ancestral functions proposed for LFY within the vascular plants (Theissen and Melzer, 2007; Moyroud et al., 2010) cannot be resolved without functional validation. Here we present a functional analysis in Ceratopteris that reveals a stem cell maintenance role for *LFY* in both gametophyte and sporophyte shoots and discuss how that role informs our mechanistic understanding of developmental innovations during land plant evolution.

## RESULTS

### The *CrLFY1* and *CrLFY2* genes duplicated recently within the fern lineage

The *LFY* gene family is present as a single gene copy in most land plant genomes (Sayou et al., 2014). In this regard, the presence of two *LFY* genes in Ceratopteris (Himi et al., 2001) is atypical. To determine whether this gene duplication is more broadly represented within the ferns and related species (hereafter ‘ferns’), a previous amino acid alignment of LFY orthologs (Sayou et al., 2014) was pruned and supplemented with newly-available fern homologs (see Materials and Methods) to create a dataset of 120 sequences, ~50% of which were from the fern lineage (**Supplementary Files 1-3**). The phylogenetic topology inferred within the vascular plants using the entire dataset (**Figure 1-figure supplement 1**) was consistent with previous analyses (Qiu et al., 2006; Wickett et al., 2014). Within the ferns (64 in total), phylogenetic relationships between *LFY* sequences indicated that the two gene copies identified in *Equisetum arvense, Azolla caroliniana* and Ceratopteris each resulted from recent independent duplication events (**Figure 1**). Gel blot analysis confirmed the presence of no more than two *LFY* genes in the Ceratopteris genome (**Figure 1-figure supplement 2**). Given that the topology of the tree excludes the possibility of a gene duplication prior to diversification of the ferns, *CrLFY1* and *CrLFY2* are equally orthologous to the single copy *LFY* representatives in other fern species.

**Figure 1.**
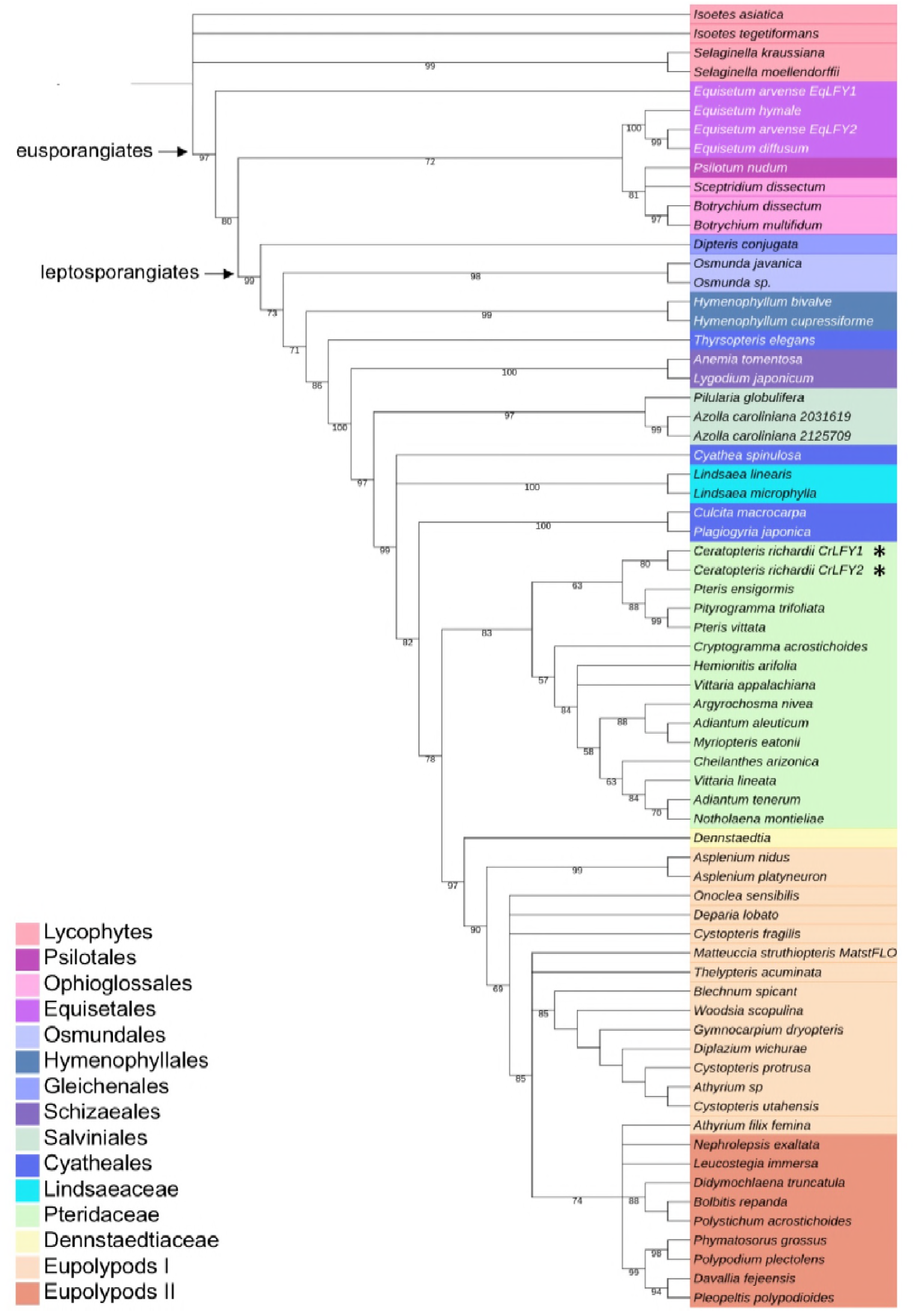
*CrLFY1* and *CrLFY2* arose from a recent gene duplication event. Inferred phylogenetic tree from maximum likelihood analysis of 64 LFY amino acid sequences (see **Supplementary File 1** for accession numbers) sampled from within the fern lineage plus lycophyte sequences as an outgroup. Bootstrap values are given for each node. The tree shown is extracted from a phylogeny with representative sequences from all land plant lineages (**Figure 1-figure supplement 1**). The *Ceratopteris richardii* genome contains no more than two copies of LFY (**Figure 1-figure supplement 2**; indicated by **\***). Different taxonomic clades within the fern lineage are denoted by different colours, as shown. The divergence between eusporangiate and leptosporangiate ferns is indicated by arrows.

### *CrLFY1* and *CrLFY2* transcripts accumulate differentially during the Ceratopteris lifecycle

The presence of two *LFY* genes in the Ceratopteris genome raises the possibility that gene activity was neo- or sub-functionalized following duplication. To test this hypothesis, transcript accumulation patterns of *CrLFY1* and *CrLFY2* were investigated throughout the Ceratopteris lifecycle. The sampled developmental stages spanned from imbibed spores prior to germination (**Figure 2A**), to differentiated male and hermaphrodite gametophytes (**Figure 2B-D**), through fertilization and formation of the diploid embryo (**Figure 2E**), to development of the increasingly complex sporophyte body plan (**Figure 2F-K**). Quantitative RT-PCR analysis detected transcripts of both *CrLFY1* and *CrLFY2* at all stages after spore germination, but only *CrLFY2* transcripts were detected in spores prior to germination (**Figure 2L**). A two-way ANOVA yielded a highly significant interaction (F(10,22) = 14.21; p < 0.0001) between gene copy and developmental stage that had not been reported in earlier studies (Himi et al., 2001), and is indicative of differential *CrLFY1/2* gene expression that is dependent on developmental stage. Of particular note were significant differences between *CrLFY1* and *CrLFY2* transcript levels during sporophyte development. Whereas *CrLFY2* transcript levels were comparable across sporophyte samples, *CrLFY1* transcript levels were much higher in samples that contained the shoot apex (**Figure 2F**, G) than in those that contained just fronds (**Figure 2H-K**). These data suggest that *CrLFY1* and *CrLFY2* genes may play divergent roles during sporophyte development, with *CrLFY1* acting primarily in the shoot apex and *CrLFY2* acting more generally.

**Figure 2.**
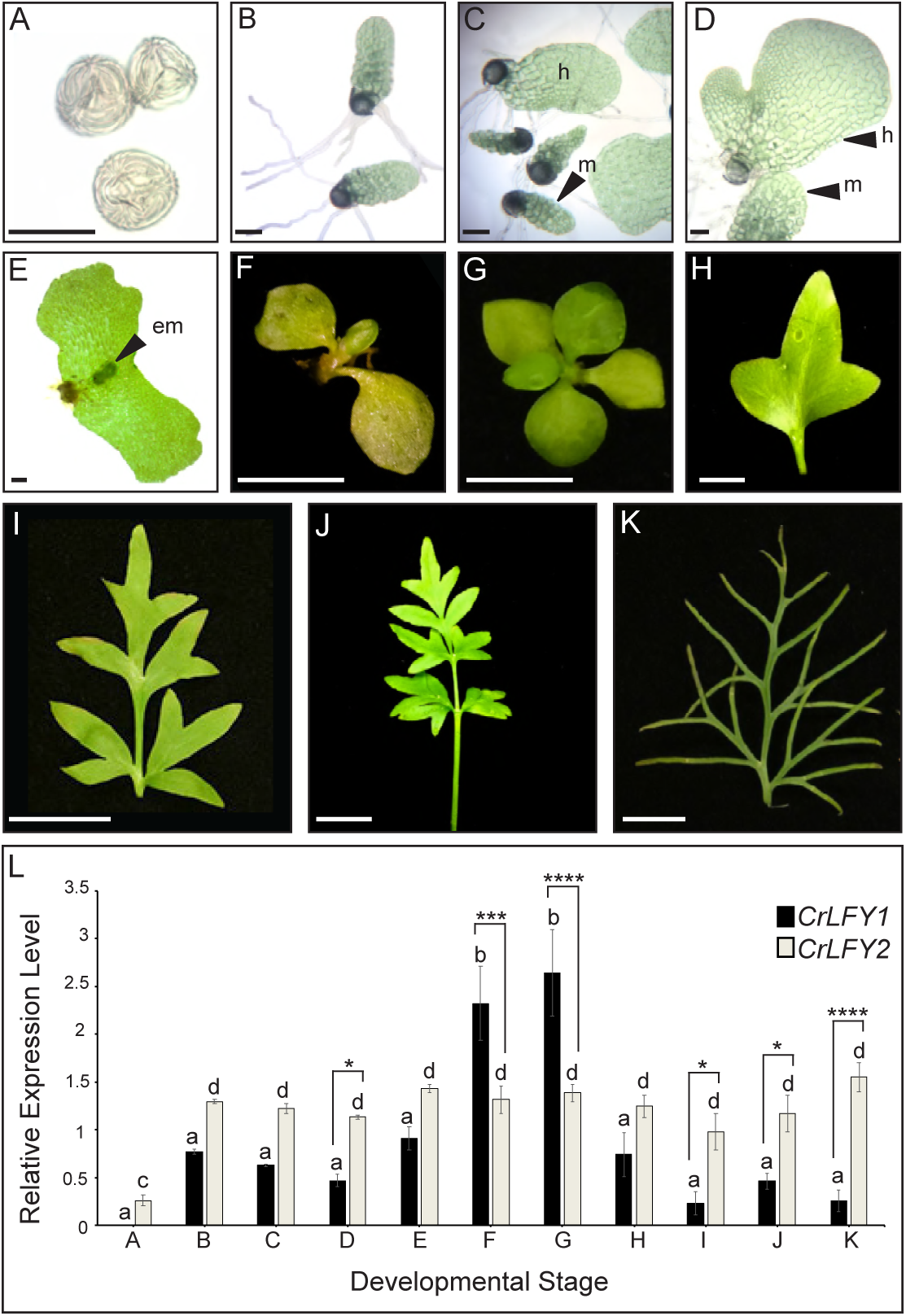
*CrLFY1* and *CrLFY2* are differentially expressed during the Ceratopteris lifecycle. **A-K**. Representative images of the developmental stages sampled for expression analysis in (**L**). Imbibed spores (**A**); populations of developing gametophytes harvested at 5 (**B**, **C**) and 8 (**D**) days after spore-sowing (DPS), comprising only males (**B**) or a mixture of hermaphrodites (h) and males (m) (**C, D**); fertilized gametophyte subtending a developing sporophyte embryo (em) (**E**); whole sporophyte shoots comprising the shoot apex with 3 (**F**) or 5 expanded entire fronds attached (**G**); individual vegetative fronds demonstrating a heteroblastic progression in which frond complexity increases through successive iterations of lateral outgrowths (pinnae) (**H**-**J**); complex fertile frond with sporangia on the underside of individual pinnae (**K**). Scale bars = 100 um (**A-E**), 5 mm (**F-H**), 20 mm (**I-K**). **L**. Relative expression levels of *CrLFY1* and *CrLFY2* (normalised against the housekeeping genes *CrACTIN1* and *CrTBP*) at different stages of development. *n* = 3; Error bars = standard error of the mean (SEM). Pairwise statistical comparisons (ANOVA followed by Tukey’s multiple comparisons test–**Supplementary File 4**) found no significant difference in *CrLFY2* transcript levels between any gametophyte or sporophyte tissues sampled after spore germination (p > 0.05) and no significant difference between *CrLFY1* and *CrLFY2* transcript levels during early gametophyte development (p>0.05) (**B, C**). Differences between *CrLFY1* and *CrLFY2* transcript levels were significant in gametophytes at 8 DPS (P<0.05) (**D**). *CrLFY1* transcript levels were significantly higher in whole young sporophytes (**F**) and vegetative shoots (**G**) compared to isolated fronds (**H-K**) (p < 0.05). *CrLFY1* transcript levels in whole sporophytes and shoots were greater than *CrLFY2*, whereas in isolated fronds *CrLFY1* transcript levels were consistently lower than *CrLFY2* (p < 0.05). Asterisks denote significant difference (*, p < 0.05; **, p < 0.01, ***, p < 0.001; ****, p < 0.0001) between *CrLFY1* and *CrLFY2* transcript levels (Sidak’s multiple comparisons test) within a developmental stage. Letters denote significant difference (p < 0.05) between developmental stages for *CrLFY1* or *CrLFY2* (Tukey’s test). Groups marked with the same letter are not significantly different from each other (p > 0.05). Statistical comparisons between developmental stages were considered separately for *CrLFY1* and *CrLFY2*. The use of different letters between *CrLFY1* and *CrLFY2* does not indicate a significant difference.

### Spatial expression patterns of *CrLFY1* are consistent with a retained ancestral role to facilitate cell divisions during embryogenesis

Functional characterization in *P. patens* previously demonstrated that LFY promotes cell divisions during early sporophyte development (Tanahashi et al., 2005). To determine whether the spatial domains of *CrLFY1* expression are consistent with a similar role in Ceratopteris embryo development, transgenic lines were generated that expressed the reporter gene B-glucuronidase (GUS) driven by 3.9 kb of the *CrLFY1* promoter (*CrLFY1_pro_::GUS*) (**Supplementary File 5**). GUS activity was monitored in individuals from three independent transgenic lines, sampling both before and up to six days after fertilization (**Figure 3A-O**), using wild-type individuals as negative controls (**Figure 3P-T**) and individuals from a transgenic line expressing GUS driven by the constitutive 35S promoter (*35S_pro_*) (**Supplementary File 5**) as positive controls (**Figure 3U-Y**). Notably, no GUS activity was detected in unfertilized archegonia of *CrLFY1_pro_::GUS* gametophytes (**Figure 3A, F, K**) but by two days after fertilization (DAF) GUS activity was detected in most cells of the early sporophyte embryo (**Figure 3B, G, L**). At 4 DAF, activity was similarly detected in all visible embryo cells, including the embryonic frond, but not in the surrounding gametophytic tissue (the calyptra) (**Figure 3C, H, M**). This embryo-wide pattern of GUS activity became restricted in the final stages of development such that by the end of embryogenesis (6 DAF) GUS activity was predominantly localized in the newly-initiated shoot apex (**Figure 3D, E, I, J, N, O**). Collectively, the GUS activity profiles indicate that *CrLFY1* expression is induced following formation of the zygote, sustained in cells of the embryo that are actively dividing, and then restricted to the shoot apex at embryo maturity. This profile is consistent with the suggestion that *CrLFY1* has retained the LFY role first identified in *P. patens* (Tanahashi et al., 2005), namely to promote the development of a multicellular sporophyte, in part by facilitating the first cell division of the zygote.

**Figure 3.**
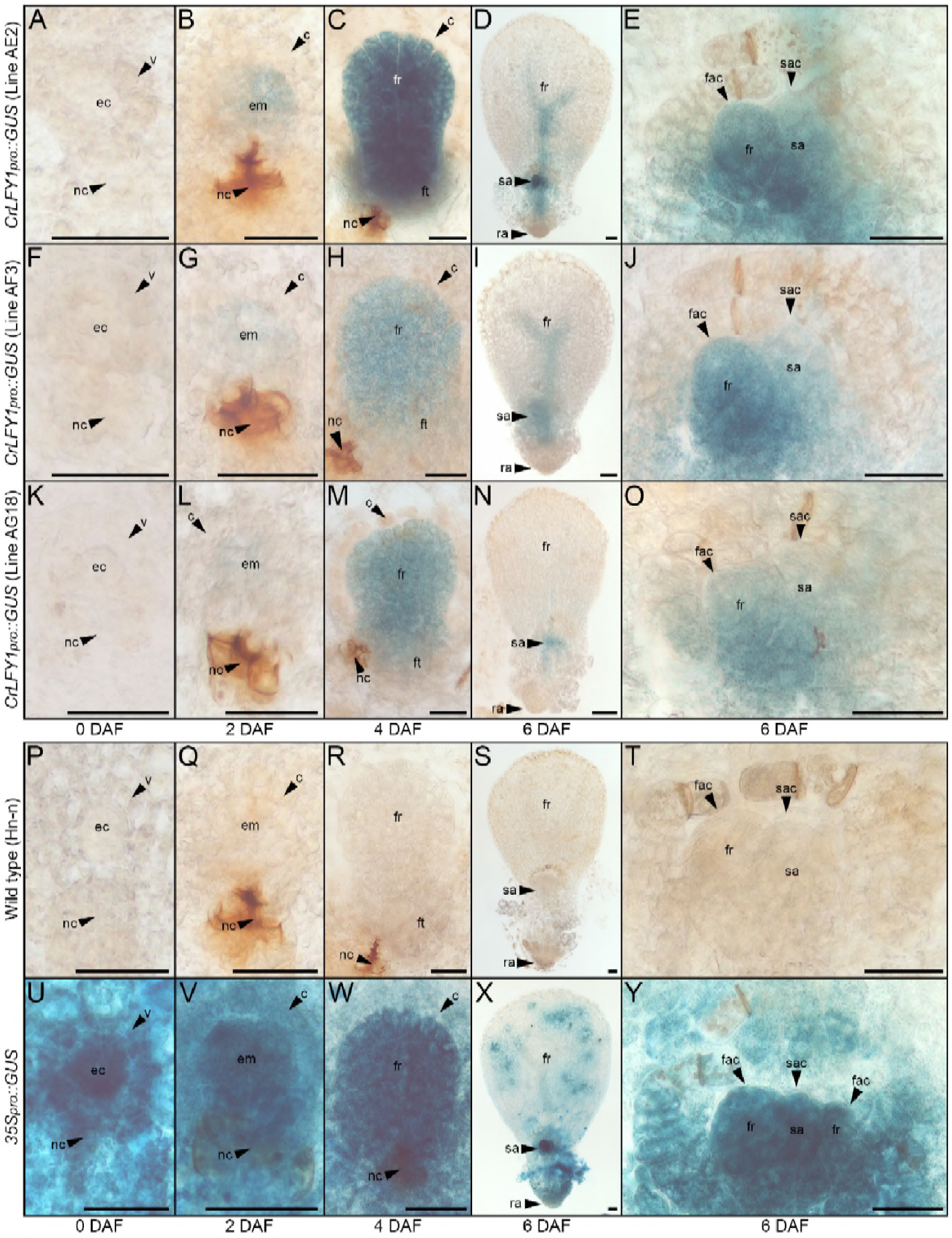
The *CrLFY1* promoter drives reporter gene expression in proliferating tissues of the developing Ceratopteris embryo. **A-Y**. GUS activity detected as blue staining in developing embryos of three independent *CrLFY1_pro_::GUS* transgenic reporter lines (**A-O**), a representative negative wild-type control line (**P-T**) and a representative positive *35S_pro_::GUS* control line (**U-Y**). Tissues are shown prior to fertilization (**A, F, K, P, U**), or 2 (**B, G, L, Q, V**), 4 (**C, H, M, R, W**), and 6 (**D, I, N, S, X**) days after fertilization (DAF). In *CrLFY1_pro_::GUS* lines, GUS activity first became visible within the first few divisions of embryo development (but not in surrounding gametophyte tissues) at 2 DAF (**B, G, L**) and was expressed in cells of the embryo frond as it proliferated (**C, H, M**). GUS activity was visible in the shoot apex at 6 DAF (**D, I, N**), with staining in the shoot apical cell (sac), subtending shoot apex tissues and newly-initiated fronds, including the frond apical cell (fac) (**E, J, O**). No GUS activity was detected in wild-type samples (**P-T**), whereas the majority of cells in the constitutively expressing *35S_pro_::GUS* samples stained blue (**U-Y**). Embryos develop on the surface of the gametophyte thallus when an egg cell (ec) within the archegonium (which comprises a venter (v) and neck canal (nc) to allow sperm entry) are fertilized. After fertilization, the venter forms a jacket of haploid cells known as the calyptra (c) that surrounds the diploid embryo (em). Cell fates in the embryo (embryo frond (fr), embryo foot (ft), root apex (ra) and shoot apex (sa)) are established at the eight-celled stage (Johnson & Renzaglia, 2008), which is around 2 DAF under our growth conditions. Embryogenesis is complete at 6 DAF, after which fronds arise from the shoot apex. Scale bars = 50 μm.

### *CrLFY1* is expressed in dividing tissues throughout shoot development

Both mosses and ferns form embryos, but moss sporophyte development is determinate post-embryogenesis (Kato and Akiyama, 2005; Kofuji and Hasebe, 2014) whereas fern sporophytes are elaborated from indeterminate shoot apices (Bierhorst, 1977; White and Turner, 1995). *CrLFY1* expression in the shoot apex at the end of embryogenesis (**Figure 3E, J, O**) and elevated transcript levels in shoot apex-containing sporophyte tissues (**Figure 2L**) suggested an additional role for *CrLFY1* relative to that seen in mosses, namely to promote proliferation in the indeterminate shoot apex. To monitor *CrLFY1* expression patterns in post-embryonic sporophytes, GUS activity was assessed in *CrLFY1_pro_::GUS* lines at two stages of vegetative development (**Figure 4A-O**) and after the transition to reproductive frond formation (**Figure 4-figure supplement 1**A-L). Wild-type individuals were used as negative controls (**Figure 4P-T**; **Figure 4-figure supplement 1**M-P) and *35S_pro_::GUS* individuals as positive controls (**Figure 4U-Y**; **Figure 4-figure supplement 1Q-T**). In young sporophytes (20 DAF), GUS activity was primarily localized in shoot apical tissues and newly-emerging frond primordia (**Figure 4A, F, K**), with very little activity detected in the expanded simple fronds (**Figure 4B, G, L**). In older vegetative sporophytes (60 DAF), which develop complex dissected fronds (**Figure 4C, H, M**), GUS activity was similarly localized in the shoot apex and young frond primordia in two out of the three fully characterized lines (**Figure 4D, I, N**) and in a total of 8 out of 11 lines screened (from seven independent rounds of plant transformation). GUS activity was also detected in developing fronds in regions where the lamina was dividing to generate pinnae and pinnules (**Figure 4E, J, O**). In some individuals GUS activity could be detected in frond tissues almost until maturity (**Figure 4C**). Notably, patterns of *CrLFY1_pro_::GUS* expression were the same in the apex and complex fronds of shoots before (60 DAF) (**Figure 4C-E, H-J, M-O**) and after (~ 115 DAF) the reproductive transition (**Figure 4-figure supplement 1A-L**). Consistent with a general role for *CrLFY1* in promoting cell proliferation in the shoot, GUS activity was also detected in shoot apices that initiate *de novo* at the lamina margin between pinnae (**Figure 4Z-AD**). Together these data support the hypothesis that *LFY* function was recruited to regulate cell division processes in the shoot when sporophytes evolved from determinate to indeterminate structures.

**Figure 4.**
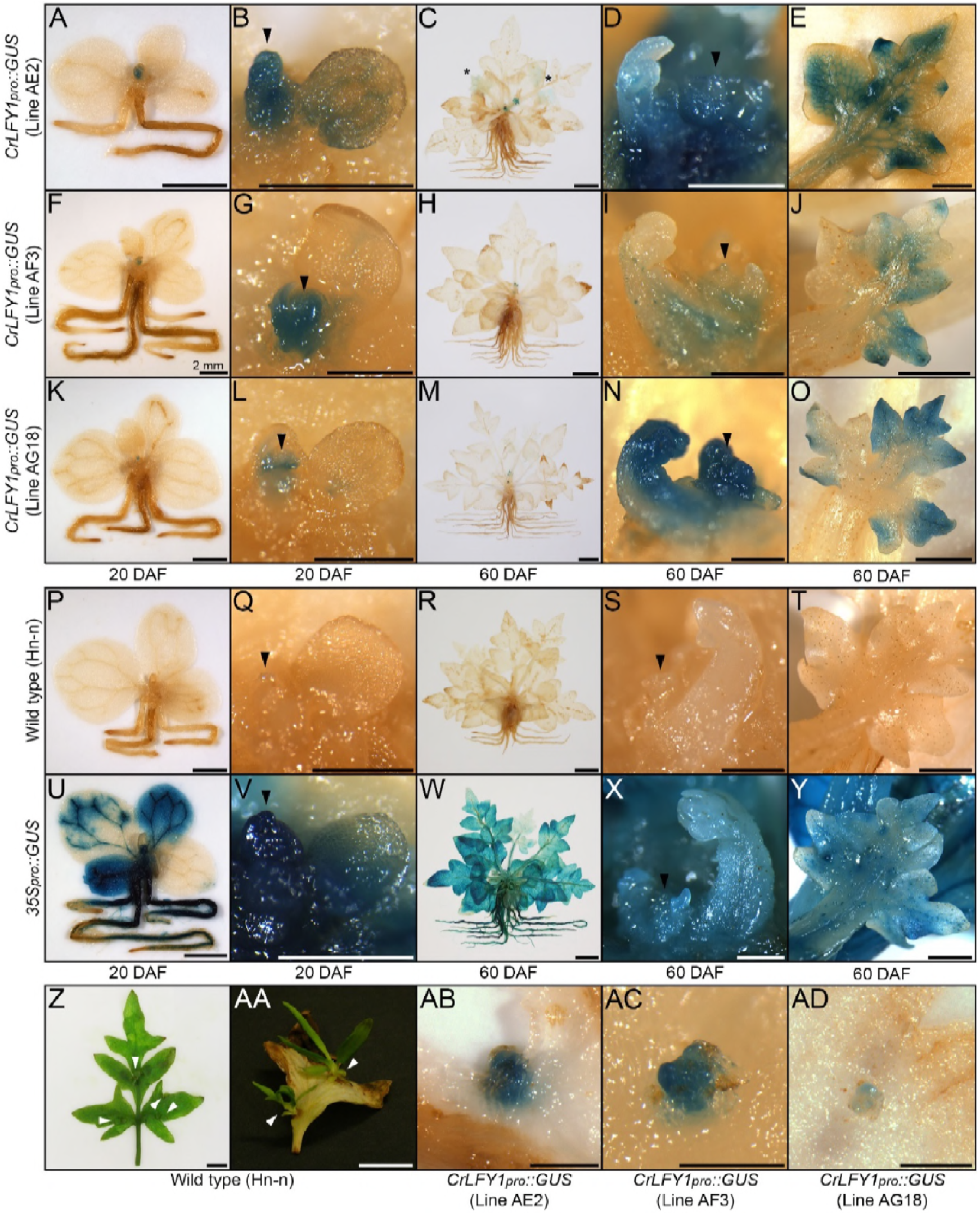
The *CrLFY1* promoter drives reporter gene expression in proliferating shoot tissues of the Ceratopteris sporophyte. **A-Y**. GUS activity detected as blue staining in post-embryonic sporophytes from three independent *CrLFY1_pro_::GUS* transgenic reporter lines (**A-O**), negative wild-type controls (**P-T**) and positive *35S_pro_::GUS* controls (**U-Y**). Sporophytes were examined at 20 DAF (**A, B, F, G, K, L, P, Q, U, V**) and 60 DAF (**C-E**, **H-J**, **M-O**, **R-T**, **W-Y**). GUS staining patterns are shown for whole sporophytes (**A, C, F, H, K, M, P, R, U, W**), shoot apices (black arrowheads) (**B, D, G, I, L, N, Q, S, V, X**) and developing fronds (**E, J, O, T, Y**). In *CrLFY1_pro_::GUS* sporophytes at 20 DAF (producing simple, spade-like fronds) GUS activity was restricted to the shoot apex (**A, F, K**) and newly-initiated frond primordia, with very low activity in expanded fronds (**B, G, L)**. In *CrLFY1_pro_::GUS* sporophytes at 60 DAF (producing complex, highly dissected fronds) GUS activity was similarly restricted to the apex (**C, H, M**), but persisted for longer during frond development. Activity was initially detected throughout the frond primordium (**D, I, N**), before becoming restricted to actively proliferating areas of the lamina (**E, J, O**). Scale bars = 2 mm (**A, F, K, P, U**), 500 μm (**B, D, G, I, L, N, Q, S, V, X**) 10 mm (**C, H, M, R, W**), 1 mm (**E, J, O, T, Y**). * = GUS staining in maturing frond. GUS staining patterns were the same in leaves formed after the reproductive transition (**Figure 4-figure supplement 1**). **Z-AD**. Fronds can initiate *de novo* shoots (white arrowheads) from marginal tissue between existing frond pinnae (**Z**, **AA**). GUS activity was detected in emerging *de novo* shoot apices on *CrLFY1_pro_::GUS* fronds (**AB-AD**). Scale bars = 10 mm (**Z, AA**), 500 μm (**AB-AD**).

### *CrLFY1* regulates activity of the sporophyte shoot apex

To test the functional significance of *CrLFY* expression patterns, transgenic RNAi lines were generated in which one of four RNAi constructs targeted to *CrLFY1, CrLFY2* or both were expressed from the maize ubiquitin promoter (*ZmUbi_pro_*) (**Supplementary File 6**). Phenotypic screening identified 10 lines with similar developmental defects that were associated with reduced *CrLFY* expression (Table 1).

**Table 1.** Summary of *CrLFY* RNAi transgenic lines and their phenotypic characterization. Transgenic lines exhibited gametophytic developmental arrest and/or sporophyte shoot termination at varying stages of development. ‘+’ indicates that a particular line was phenotypically normal at the developmental stage indicated, ‘−’ indicates that development had arrested at or prior to this stage. In lines marked ‘+/−’ the stage at which developmental defects occurred varied between individuals within the line, and at least some arrested individuals were identified at the stage indicated. The five *ZmUbi_pro_::CrLFY1/2-i1* lines shown were generated from three rounds of transformation, the pairs of lines B16 and B19 and D2 and D4 potentially arising from the same transformation event. For all other constructs each transgenic line arose from a separate round of transformation and so must represent independent T-DNA insertions.

| RNAi transgene | Line | Transformation replicate | Gametophyte phase |  |  |  | Sporophyte phase |  |  |  |  |
| --- | --- | --- | --- | --- | --- | --- | --- | --- | --- | --- | --- |
|  |  |  | Spore germination | AC-based growth | Notch meristem-based growth | % Population arrested | Embryo | Shoot apex initiated | Simple frond | Complex frond | % Population arrested |
| *ZmUbi<sub>pro</sub>::CrLFY1/2-i1* | B16 | 1 | + | - | - | 99.86 | + | + | - | - | <5% |
| *ZmUbi<sub>pro</sub>::CrLFY1/2-i1* | B19 | 1 | + | - | - | 50.00 | + | + | + | - | <5% |
| *ZmUbi<sub>pro</sub>::CrLFY1/2-i1* | D13 | 2 | + | - | - | 99.80 | + | + | - | - | <5% |
| *ZmUbi<sub>pro</sub>::CrLFY1/2-i1* | D2 | 3 | + | + | + | 0.00 | + | + | - | - | <5% |
| *ZmUbi<sub>pro</sub>::CrLFY1/2-i1* | D4 | 3 | + | + | + | 0.00 | + | + | + | + | <5% |
| *ZmUbi<sub>pro</sub>::CrLFY1/2-i2* | F9 | 4 | + | - | - | 0.00 | + | + | - | - | <5% |
| *ZmUbi<sub>pro</sub>::CrLFY1/2-i2* | F14 | 5 | - | - | - | 100.00 | - | - | - | - | - |
| *ZmUbi<sub>pro</sub>::CrLFY1-i3* | E8 | 6 | + | +/- | +/- | 100.00 | - | - | - | - | - |
| *ZmUbi<sub>pro</sub>::CrLFY1-i3* | G13 | 7 | + | + | + | 0.00 | + | + | - | - | <5% |
| *ZmUbi<sub>pro</sub>::CrLFY2-i4* | C3 | 9 | + | + | + | 0.00 | + | + | - | - | <5% |

Wild-type post-embryonic shoot development begins with the production of simple, spade-like fronds from the shoot apex (**Figure 5A**). In eight transgenic lines, sub-populations of sporophytes developed in which early sporophyte development was perturbed, one line (E8) failing to initiate recognizable embryos (**Figure 5B**) and the remainder exhibiting premature shoot apex termination, typically after producing several distorted fronds (**Figure 5C-H**). Sub-populations of phenotypically normal transgenic sporophytes were also identified in some of these lines (**Figure 5I-L**). The two remaining lines exhibited less severe shoot phenotypes, one (B19) undergoing shoot termination only when wild-type sporophytes produced complex fronds (**Figure 5M, N**) and the other (D4a) completing sporophyte development but reduced in size to approximately 50% of wild-type (**Figure 5O, P**). Despite the predicted sequence specificity of *CrLFY1-i3* and *CrLFY2-i4* (**Supplementary File 6**), quantitative RT-PCR analysis found that all four RNAi constructs led to suppressed transcript levels of both *CrLFY* genes (**Figure 5Q**). The severity of the shoot phenotype was correlated with the level of endogenous *CrLFY* transcripts detected across all lines (**Figure 5Q**), with relative levels of both *CrLFY1* and *CrLFY2* significantly reduced compared to wild-type in all early-terminating sporophytes (p < 0.0001 and p < 0.01, respectively). Notably, phenotypic differences between the two less-severe lines correlated with differences in *CrLFY1* transcript levels, which were significantly reduced (p < 0.01) in late-terminating line B19 but not significantly different from wild-type (p > 0.05) in the non-arresting line D4a. *CrLFY2* expression was not significantly different from wild-type (p > 0.05) in either the B19 or D4a line, or in the phenotypically normal transgenics. Together these data indicate that wild-type levels of *CrLFY2* are sufficient to compensate for some loss of *CrLFY1*, but at least 10% of wild-type *CrLFY1* activity is required to prevent premature termination of the shoot apex. It can thus be concluded that *CrLFY1* and *CrLFY2* act partially redundantly to maintain indeterminacy of the shoot apex in Ceratopteris, a role not found in the early divergent bryophyte *P. patens*, nor known to be retained in the majority of later diverging flowering plants.

**Figure 5.**
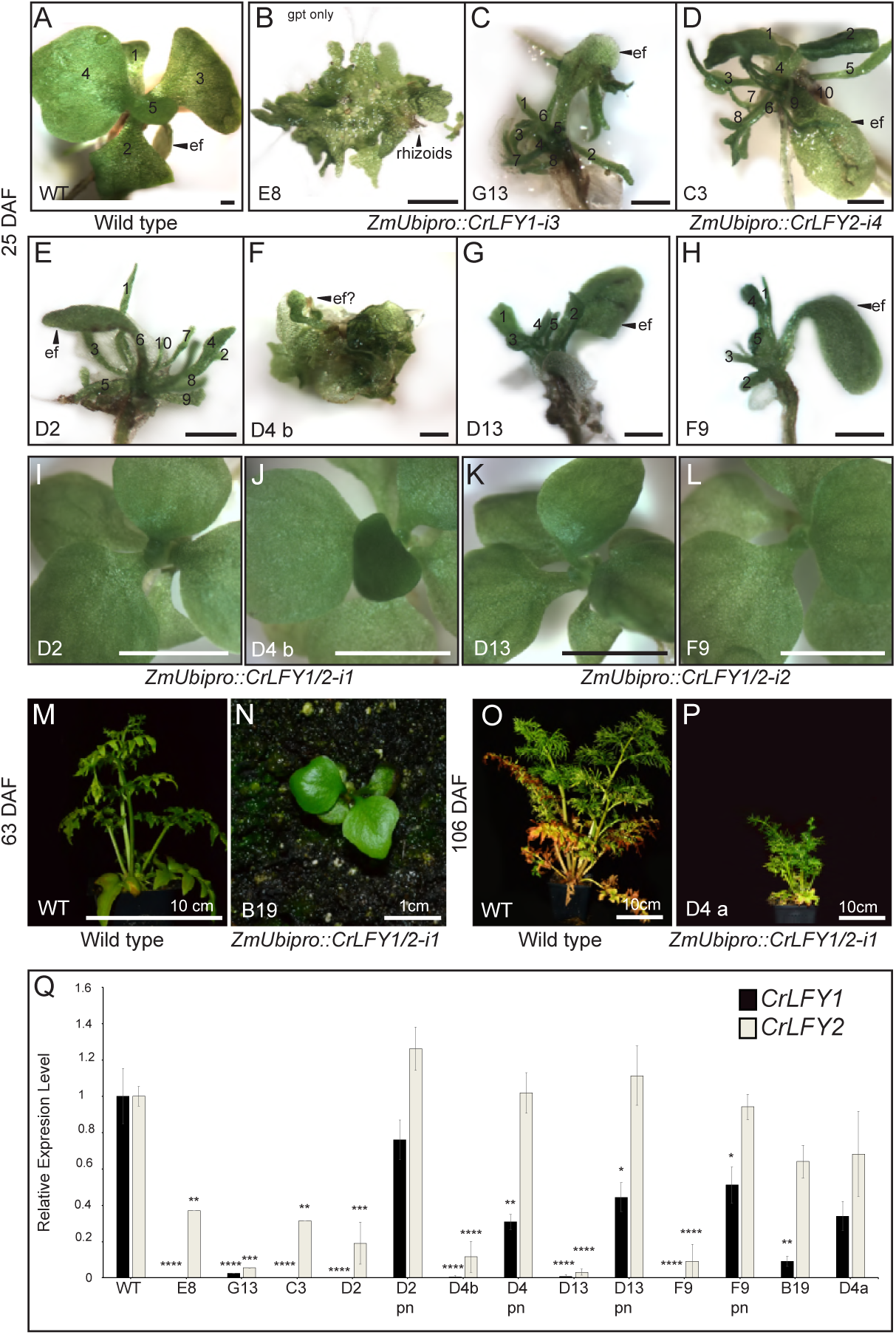
Suppression of *CrLFY* expression causes early termination of the Ceratopteris sporophyte shoot apex. **A-L**. Sporophyte phenotype 25 days after fertilization (DAF) in wild-type (**A**) and transgenic lines carrying RNAi constructs against *CrLFY1* (*ZmUbi_pro_::CrLFY1-i3*) (**B**, **C**), *CrLFY2* (*ZmUbi_pro_::CrLFY1-i4*) (**D**) and both *CrLFY1* and *CrLFY2* (*ZmUbi_pro_::CrLFY1/2-i1* and *ZmUbi_pro_::CrLFY1/2-i2*) (**E-L**). In some lines, both aborted and phenotypically normal sporophytes were identified (compare **E** & **I**; **F** & **J**; **G** & **K**; **H** & **L**). The presence of the RNAi transgene in phenotypically normal sporophytes was validated by genotyping (**Supplementary File 6**). Scale bars = 100 μm (**A-H**), 5mm (**I-L**). **M-P**. Sporophyte phenotype of wild-type (**M, O**) and two *ZmUbi_pro_::CrLFY1/2-i1* lines (**N, P**) at 63 (**M, N)**) and 106 (**O, P**) DAF. Scale bars =1cm (**O**), 10cm (**M, O, P**). **Q**. qRT-PCR analysis of *CrLFY1* and *CrLFY2* transcript levels (normalized against the averaged expression of housekeeping genes *CrACTIN1* and *CrTBP*) in the sporophytes of the RNAi lines shown in (**A-P**). Transcript levels are depicted relative to wild type. *n* = 3, error bars = standard error of the mean (SEM). *CrLFY1* and *CrLFY2* expression levels were significantly reduced compared to wild type (≤1% and 3-19%, respectively) in all transgenic lines where sporophyte shoots undergo early termination (**A-H**), but in phenotypically normal (pn) sporophytes segregating in the same lines (**I-L**), only *CrLFY1* transcript levels were reduced (D4b, p < 0.01; D13, F9, p < 0.05). *CrLFY2* transcript levels were not significantly different from wild-type in the phenotypically normal sporophytes at 25 DAF (**I-L**) or in the sporophytes that survived to 63 (**N**) or 106 (**P**) DAF. *CrLFY1* levels were significantly reduced (~10%) in line B19 where shoot termination occurred ~63 DAF (**N**) but were not significantly different from wild type in line D4a where shoot development did not terminate prematurely (**P**). Asterisks denote level of P were significant difference (*, p < 0.05; **, p < 0.01, ***, p < 0.001; ****, p < 0.0001) from wild type.

### *CrLFY* promotes apical cell divisions in the gametophyte

In six of the RNAi lines that exhibited sporophyte developmental defects, it was notable that 50%-99% of gametophytes arrested development prior to the sporophyte phase of the lifecycle (Table 1). This observation suggested that LFY plays a role in Ceratopteris gametophyte development, a function not previously recognized in either bryophytes or angiosperms. During wild-type development, the Ceratopteris gametophyte germinates from a single-celled haploid spore, establishing a single apical cell (AC) within the first few cell divisions (**Figure 6A**). Divisions of the AC go on to form a two-dimensional photosynthetic thallus in both the hermaphrodite (**Figure 6B**) and male sexes (**Figure 6C**). In contrast, the gametophytes from six RNAi lines (carrying either *ZmUbi_pro_::CrLFY1-i3, ZmUbi_pro_::CrLFY1/2-i1* or *ZmUbi_pro_::CrLFY1/2-i2*) exhibited developmental arrest (**Figure 6D-J**), which in five lines clearly related to a failure of AC activity. The point at which AC arrest occurred varied, in the most severe line occurring prior to or during AC specification (**Figure 6D**) and in others during AC-driven thallus proliferation (**Figure 6E-I**). Failure of AC activity was observed in both hermaphrodites (**Figure 6E**) and males (**Figure 6H, I**). The phenotypically least severe line exhibited hermaphrodite developmental arrest only after AC activity had been replaced by the notch meristem (**Figure 6J**). A role for *CrLFY* in maintenance of gametophyte AC activity was supported by the detection of *CrLFY* transcripts in the AC and immediate daughter cells of wild-type gametophytes (**Figure 6K-N**). By contrast *CrLFY* transcripts were not detected in arrested *ZmUbi_pro_::CrLFY1/2-i1* lines (**Figure 6O-R**) despite confirmed presence of the transgene (**Figure 6-figure supplement 1**). Although the observed phenotypes could not be ascribed to a specific gene copy, these data support a role for *CrLFY* in AC maintenance during gametophyte development, and thus invoke a role for LFY in the regulation of apical activity in both the sporophyte and gametophyte phases of vascular plant development.

**Figure 6.**
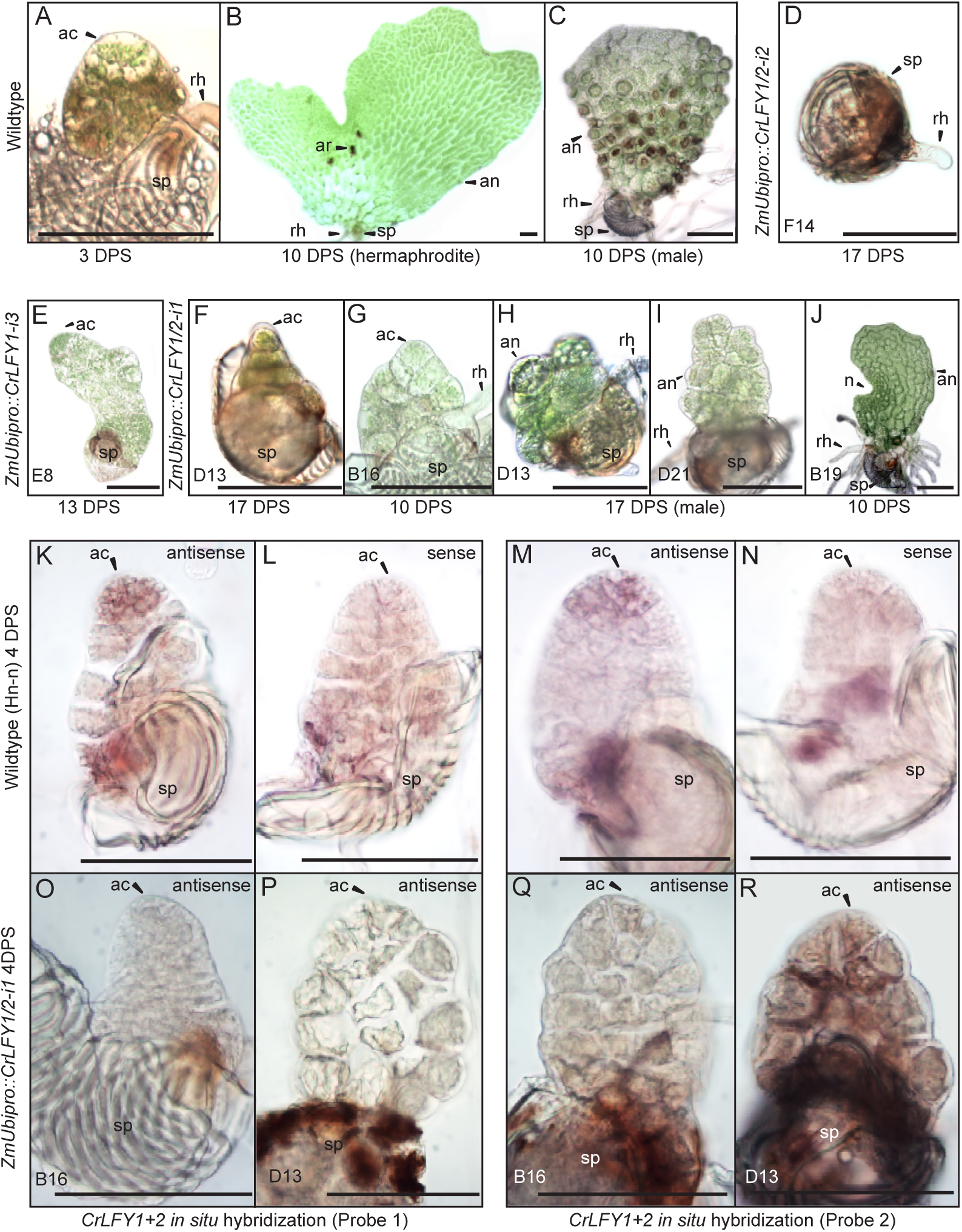
Suppression of *CrLFY* expression causes early termination of the Ceratopteris gametophyte apical cell. **A-C**. The wild-type Ceratopteris gametophyte establishes a triangular apical cell (ac) shortly after spore (sp) germination (**A**). Divisions of the apical cell establish a photosynthetic thallus in both hermaphrodite and male gametophytes. At 10 days post spore sowing (DPS) both gametophyte sexes are approaching maturity, with the hermaphrodite (**B**) having formed a chordate shape from divisions at a lateral notch meristem (n) and having produced egg-containing archegonia (ar), sperm-containing antheridia (an), and rhizoids (rh). The male (**C**) has a more uniform shape with antheridia across the surface. **D-J**. When screened at 10-17 DPS, gametophytes from multiple RNAi lines (as indicated) exhibited developmental arrest, mostly associated with a failure of apical cell activity. Arrest occurred at various stages of development from failure to specify an apical cell, resulting in only a rhizoid being produced and no thallus (**D**) through subsequent thallus proliferation (**E-I**). Gametophyte development in one line progressed to initiation of the notch meristem but overall thallus size was severely reduced compared to wild-type (**J**). **K-R**. *In situ* hybridization with antisense probes detected *CrLFY* transcripts in the apical cell and immediate daughter cells of wild-type gametophytes at 4 DPS (**K, M**). No corresponding signal was detected in controls hybridized with sense probes (**L, N**). In the arrested gametophytes of two *ZmUbi_pro_*::*CrLFY1/2-i1* lines *CrLFY* transcripts could not be detected (**O-R**), and transgene presence was confirmed (**Figure 6-figure supplement 1**). Scale bars = 100 μm.

## DISCUSSION

The results reported here reveal a role for LFY in the maintenance of apical cell activity throughout gametophyte and sporophyte shoot development in Ceratopteris. During sporophyte development, qRT-PCR and transgenic reporter lines demonstrated that *CrLFY1* is preferentially expressed in the shoot apex (whether formed during embryogenesis or *de novo* on fronds, and both before and after the reproductive transition); in emerging lateral organ (frond) primordia; and in pinnae and pinnules as they form on dissected fronds (**Figures 2-4**). Notably, active cell division is the main feature in all of these contexts. *CrLFY2* transcript levels were more uniform throughout sporophyte shoot development, in both dividing tissues and expanded fronds (**Figure 2**), and expression has previously been reported in roots (Himi et al., 2001). Simultaneous suppression of *CrLFY1* and *CrLFY2* activity by RNAi resulted in developmental arrest of both gametophyte and sporophyte shoot apices, with any fronds produced before termination of the sporophyte apex exhibiting abnormal morphologies (**Figures 5, 6**). The severity of phenotypic perturbations in sporophytes of transgenic lines correlated with combined *CrLFY1* and *CrLFY2* transcript levels, with wild-type levels of *CrLFY2* able to fully compensate for up to a 70% reduction in *CrLFY1* levels (**Figure 5**). The duplicate *CrLFY* genes therefore act at least partially redundantly during shoot development in Ceratopteris.

A function for LFY in gametophyte development has not previously been reported in any land plant species. In the moss *P. patens, PpLFY1* and *PpLFY2* are expressed in both the main and lateral apices of gametophytic leafy shoots but double loss of function mutants develop normally; indicating that LFY is not necessary for maintenance of apical cell activity in the gametophyte (Tanahashi et al., 2005). By contrast, loss of *CrLFY* expression from the gametophyte shoot apex results in loss of apical cell activity during thallus formation in Ceratopteris (**Figure 6**). The different DNA binding site preferences (and hence downstream target sequences) of PpLFY and CrLFY (Sayou et al., 2014) may be sufficient to explain the functional distinction in moss and fern gametophytes, but the conserved expression pattern is intriguing given that there should be no pressure to retain that pattern in *P. patens* in the absence of functional necessity. The thalloid gametophytes of the two other extant bryophyte lineages (liverworts and hornworts) resemble the fern gametophyte more closely than mosses (Ligrone et al., 2012), but LFY function in these contexts is not yet known. Overall the data are consistent with the hypothesis that in the last common ancestor of ferns and angiosperms, LFY functioned to promote cell proliferation in the thalloid gametophyte, a role that has been lost in angiosperms where gametophytes have no apical cell and are instead just few-celled determinate structures.

The range of reported roles for LFY in sporophyte development can be rationalized by invoking three sequential changes in gene function during land plant evolution (**Figure 7**). First, the ancestral LFY function to promote early cell divisions in the embryo was retained as bryophytes and vascular plants diverged, leading to conserved roles in *P. patens* (Tanahashi et al., 2005) and Ceratopteris (**Figure 3**). Second, within the vascular plants (preceding divergence of the ferns) this proliferative role expanded to maintain apical cell activity, and hence to enable indeterminate shoot growth. This is evidenced by *CrLFY* activity at the tips of shoots, fronds and pinnae (**Figures 3-5**), all of which develop from one or more apical cells (Hill, 2001; Hou and Hill, 2004). Whether fern fronds are homologous to shoots or to leaves in angiosperms is an area of debate (Tomescu, 2009; Vasco et al., 2013; Harrison and Morris, 2018), but there are angiosperm examples of LFY function in the vegetative SAM (Ahearn et al., 2001; Zhao et al., 2017), axillary meristems (Kanrar et al., 2008; Rao et al., 2008; Chahtane et al., 2013) and in actively dividing regions of compound leaves (Hofer et al., 1997; Molinero-Rosales et al., 1999; Champagne et al., 2007; Wang et al., 2008) indicating that a proliferative role in vegetative tissues has been retained in at least some angiosperm species. Consistent with the suggestion that the angiosperm floral meristem represents a modified vegetative meristem (Theißen et al., 2016), the third stage of LFY evolution would have been co-option and adaptation of this proliferation-promoting network into floral meristems, with subsequent restriction to just the flowering role in many species. This is consistent with multiple observations of *LFY* expression in both vegetative and reproductive shoots (developing cones) in gymnosperms (Mellerowicz et al.; Mouradov et al., 1998; Shindo et al., 2001; Carlsbecker et al., 2004; Vázquez-Lobo et al., 2007; Carlsbecker et al., 2013; Moyroud et al., 2017) and suggests that pre-existing *LFY*-dependent vegetative gene networks might have been co-opted during the origin of specialized sporophyte reproductive axes in ancestral seed plants, prior to the divergence of angiosperms.

**Figure 7.**
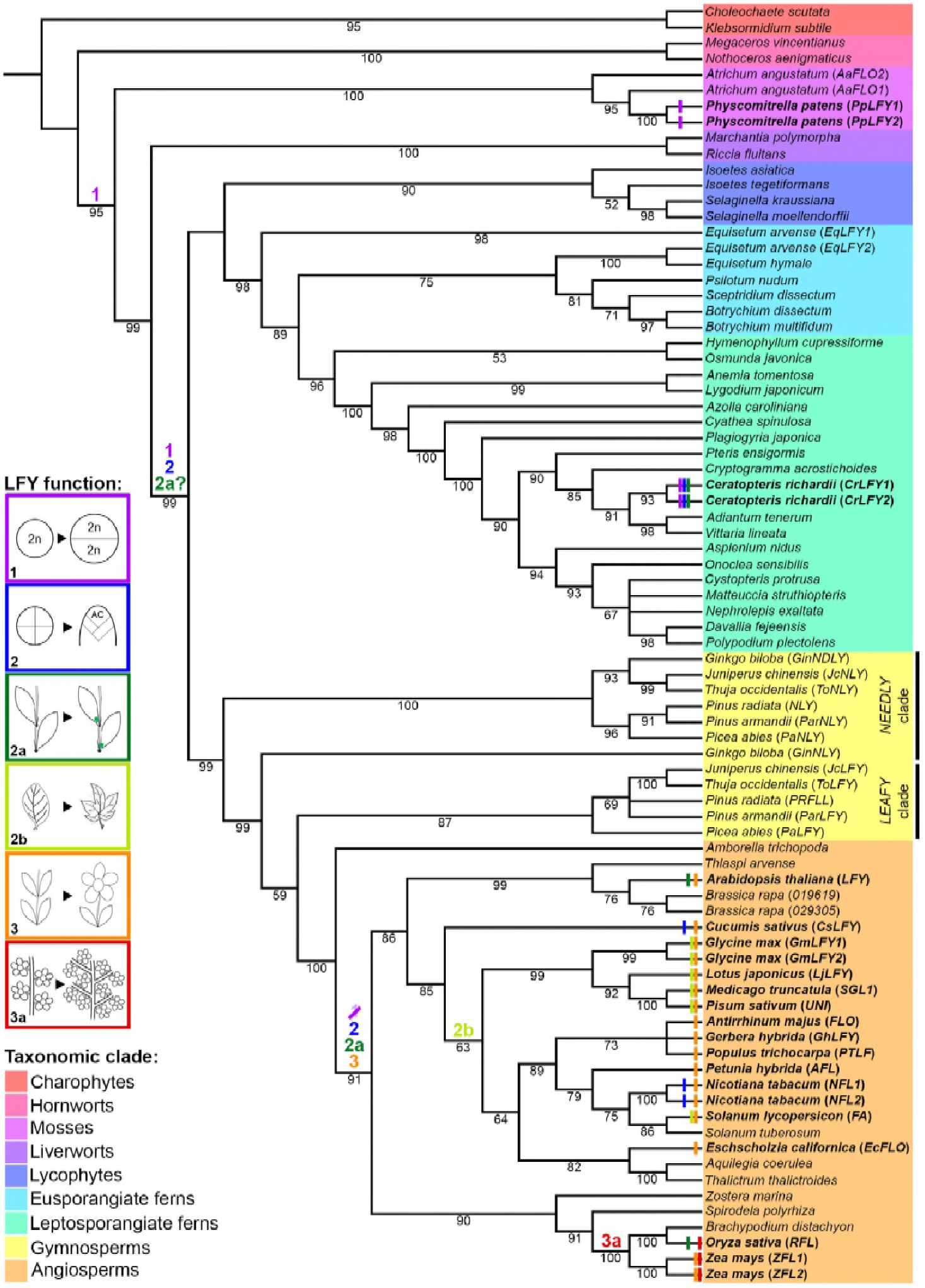
Evolutionary trajectory of LFY function. The phylogeny was reconstructed from selected LFY protein sequences representing all extant embryophyte lineages (as highlighted) and the algal sister-group. Coloured bars at the terminal branches represent different developmental functions of LFY determined from functional analysis in those species (see **Supplementary File 8** for references). Coloured numbers indicate the putative points of origin of different functions inferred from available data points across the tree. **1**, cell division within the sporophyte zygote; **2**, maintenance of indeterminate cell fate in vegetative shoots through proliferation of one or more apical cells (AC); **2a**, maintenance of indeterminate cell fate in vegetative lateral/axillary apices; **2b**, maintenance of indeterminate cell fate in the margins of developing lateral organs (compound leaves); **3**, specification of floral meristem identity (determinate shoot development producing modified lateral organs) and shoot transition to the reproductive phase; **3a**, maintenance of indeterminate cell fate in inflorescence lateral/branch meristems (in place of floral meristem fate).

The proposed evolutionary trajectory for LFY function bears some resemblance to that seen for KNOX protein function. Class I *KNOX* genes are key regulators of indeterminacy in the vegetative shoot apical meristem of angiosperms (Gaillochet et al., 2015), and are required for compound leaf formation in both tomato and *Cardamine hirsuta* (Bar and Ori, 2015). In ferns, *KNOX* gene expression is observed both in the shoot apex and developing fronds (Sano et al., 2005; Ambrose and Vasco, 2016), and in *P. patens* the genes regulate cell division patterns in the determinate sporophyte (Sakakibara et al., 2008). It can thus be speculated that *LFY* and *KNOX* had overlapping functions in the sporophyte of the last common ancestor of land plants, but by the divergence of ancestral angiosperms from gymnosperms, *KNOX* genes had come to dominate in vegetative meristems whereas *LFY* became increasingly specialized for floral meristem function. Unlike *LFY*, however, there is not yet any evidence for *KNOX* function in the gametophyte of any land plant lineage, and thus if a pathway for regulating stem cell activity was co-opted from the gametophyte into the sporophyte, it was the LFY pathway.

## MATERIALS AND METHODS

### Plant materials and growth conditions

All experimental work was conducted using *Ceratopteris richardii* strain Hn-n (Hickok et al., 1995). Plant growth conditions for Ceratopteris transformation and DNA gel blot analysis of transgenic lines were as previously described (Plackett et al., 2015).

### Phylogenetic analysis

A dataset of 99 aligned LFY protein sequences from a broad range of streptophytes was first retrieved from Sayou et al., (2014). The dataset was pruned and then supplemented with further sequences (**Supplementary File 1**) to enable trees to be inferred that would (i) provide a more balanced distribution across the major plant groups and (ii) infer fern relationships. Only a subset of available angiosperm sequences was retained (keeping both monocot and dicot representatives) but protein sequences from other angiosperm species where function has been defined through loss-of-function analyses were added from NCBI – *Antirrhinum majus* FLO AAA62574.1 (Coen *et al*. 1990), *Pisum sativum* UNI AAC49782.1 (Hofer *et al*. 1997), *Cucumis sativus* CsLFY XP_004138016.1 (Zhao *et al*. 2017), *Medicago truncatula* SGL1 AY928184 (Wang *et al*. 2008), *Petunia hybrida* ALF AAC49912.1 (Souer *et al*. 1998), *Nicotiana tabacum* NFL1 AAC48985.1 and NFL2 AAC48986.1 (Kelly et al. 1995), *Eschscholzia californica* EcFLO AAO49794.1 (Busch & Gleissberg 2003), *Gerbera hybrida* cv. ‘Terraregina’ GhLFY ANS10152.1 (Zhao *et al*. 2016), *Lotus japonicus* LjLFY AAX13294.1 (Dong *et al*. 2005) and *Populus trichocarpa* PTLF AAB51533.1 (Rottmann *et al*. 2000). To provide better resolution within and between angiosperm clades, sequences from *Spirodela polyrhiza* (32G0007500), *Zostera marina* (27g00160.1), *Aquilegia coerulea* (5G327800.1) and *Solanum tuberosum* (PGSC0003DMT400036749) were added from Phytozome v12.1 (https://phytozome.jgi.doe.gov/pz/portal.html). Genome sequence from the early-diverging Eudicot *Thalictrum thalictroides* was searched by TBLASTX (Altschul et al., 1990) (https://blast.ncbi.nlm.nih.gov/Blast.cgi?PROGRAM=tblastx&PAGE_TYPE=BlastSearch&BLAST_SPEC=&LINK_LOC=blasttab) with nucleotide sequence from the Arabidopsis *LFY* gene. A gene model was derived from sequence in two contigs (108877 & 116935) using Genewise (https://www.ebi.ac.uk/Tools/psa/genewise/). Gymnosperm sequences were retained from *Ginkgo biloba* and from a subset of conifers included in Sayou et al., (2014), whilst sequences from conifers where *in situ* hybridization patterns have been reported were added from NCBI – *Pinus radiata* PRFLL AAB51587.1 and NLY AAB68601.1 (Mellerowicz et al.; Mouradov et al., 1998) and *Picea abies* PaLFY AAV49504.1 and PaNLY AAV49503.1 (Carlsbecker et al., 2004). Fern sequences were retained except *Angiopteris spp* sequences which consistently disrupted the topology of the tree by grouping with gymnosperms. To better resolve relationships within the ferns, additional sequences were identified in both NCBI and 1KP (Matasci et al., 2014) databases. The protein sequence from *Matteuccia struthiopteris* AAF77608.1 MatstFLO (Himi et al., 2001) was retrieved from NCBI. Further sequences from horsetails (2), plus eusporangiate (1) and leptosporangiate (53) ferns were retrieved from the 1KP database (https://db.cngb.org/blast/) using BLASTP and the MatstFLO sequence as a query. Lycophyte and bryophyte sequences were all retained, but the liverwort *Marchantia polymorpha* predicted ORF sequence was updated from Phytozome v12.1 (Mpo0113s0034.1.p), the hornwort *Nothoceros* genome scaffold was replaced with a translated full length cDNA sequence (AHJ90704.1) from NCBI and two additional lycophyte sequences were added from the 1KP dataset (*Isoetes tegetiformans* scaffold 2013584 and *Selaginella kraussiana* scaffold 2008343). All of the charophyte scaffold sequences were substituted with *Coleochaete scutata* (AHJ90705.1) and *Klebsormidium subtile* (AHJ90707.1) translated full-length cDNAs from NCBI.

The new/replacement sequences were trimmed and amino acids aligned to the existing alignment from Sayou et al. (2014) using CLUSTALW (**Supplementary Files 2 and 3**). The best-fitting model parameters (JTT+I+G4) were estimated and consensus phylogenetic trees were run using Maximum Likelihood from 1000 bootstrap replicates, using IQTREE (Nguyen et al., 2015). Two trees were inferred. The first contained only a subset of fern and allied sequences to achieve a more balanced distribution across the major plant groups (81 sequences in total) (**Figure 7**), whereas the second used the entire dataset (120 sequences ~50% of which are fern and allied sequences – **Figure 1-figure supplement 1**). The data were imported into ITOL (Letunic and Bork, 2016) to generate the pictorial representations. All branches with less than 50% bootstrap support were collapsed. Relationships within the ferns (**Figure 1**) were represented by pruning the lycophyte and fern sequences (68 in total) from the tree containing all available fern sequences (**Figure 1-figure supplement 1**).

### CrLFY locus characterization and DNA gel blot analysis

Because no reference genome has yet been established for Ceratopteris (or any fern), *CrLFY* copy number was quantified by DNA gel blot analysis. Ceratopteris genomic DNA was hybridized using both the highly conserved LFY DNA-binding domain diagnostic of the *LFY* gene family (Maizel, 2005) and also gene copy-specific sequences (**Figure 1-figure supplement 2**). CrLFY1 and CrLFY2 share 85% amino acid similarity, compared to 65% and 44% similarity of each to AtLFY. DNA gel blotting and hybridization was performed as described previously (Plackett et al., 2014). The results supported the presence of only two copies of *LFY* within the Ceratopteris genome. All primers used in probe preparation are supplied in **Supplementary File 7**.

Genomic sequences for *CrLFY1* and *CrLFY2* ORFs were amplified by PCR from wild-type genomic DNA using primers designed against published transcript sequences (Himi et al., 2001). ORFs of 1551bp and 2108bp were obtained, respectively (**Figure 1-figure supplement 2**). Exon structure was determined by comparison between genomic and transcript sequences. The native promoter region of *CrLFY1* was amplified from genomic template by sequential rounds of inverse PCR with initial primer pairs designed against published *CrLFY1* 5’UTR sequence and additional primers subsequently designed against additional contiguous sequence that was retrieved. A 3.9 kb contiguous promoter fragment was isolated for *CrLFY1* containing the entire published 5’UTR and 1.9 kb of additional upstream sequence (**Figure 1-figure supplement 2**, **Supplementary File 7**).

### qPCR analysis of gene expression

RNA was extracted from Ceratopteris tissues using the Spectrum Total Plant RNA kit (Sigma-Aldrich, St. Louis, MO) and 480 ng were used as template in iScript cDNA synthesis (Bio-Rad). *CrLFY1* and *CrLFY2* locus-specific qPCR primers (**Supplementary File 7**) were designed spanning intron 1. Amplification specificity of primers was validated via PCR followed by sequencing. qPCR of three biological replicates and three technical replicates each was performed in a Bio-Rad CFX Connect with iTaq Universal SYBR Green Supermix (Bio-Rad, Hercules, CA). Primer amplification efficiency was checked with a cDNA serial dilution. Efficiency was determined using the slope of the linear regression line as calculated by Bio-Rad CFX Connect software. Primer specificity was tested via melting curve analysis, resulting in a single peak per primer set. *CrLFY* expression was calculated using the 2^−ΔΔCt^ method (Livak and Schmittgen, 2001) and normalized against the geometric mean of the expression of two endogenous housekeeping genes (Hellemans et al., 2007), *CrACTIN1* and *CrTATA-BINDING PROTEIN (TBP)* (Ganger et al., 2014). The standard deviation of the Ct values of each housekeeping gene was calculated to ensure minimal variation (<3%) in gene expression in wild-type versus transgenic lines.

Relative expression values of *CrLFY* from qPCR were compared by two-way analysis of variance (ANOVA) for developmental stages or transgenic lines, followed by Sidak’s or Tukey’s multiple comparisons in Prism v. 7.0 (GraphPad Software, Inc., La Jolla, CA). The significance threshold (p) was set at 0.05.

### Generation of GUS reporter constructs

The *CrLFY1_pro_::GUS* reporter construct (**Supplementary File 5**) was created by cloning the *CrLFY1* promoter (**Supplementary File 7**) into pART7 as a *Not*I-*Xba*I restriction fragment, replacing the existing *35S* promoter. A β-Glucuronidase (GUS) coding sequence was cloned downstream of *pCrLFY1* as an *Xba*I-*Xba*I fragment. The same GUS *Xba*I-*Xba*I fragment was also cloned into pART7 to create a *35S_pro_::GUS* positive control. The resulting *CrLFY1_pro_::GUS::ocs* and *35S_pro_::GUS::ocs* cassettes were each cloned as *Not*I-*Not*I fragments into the pART27-based binary transformation vector pBOMBER carrying a hygromycin resistance marker previously optimized for Ceratopteris transformation (Plackett et al., 2015).

### Generation of RNAi constructs

RNAi constructs were designed and constructed using the pANDA RNAi expression system (Miki and Shimamoto, 2004). Four RNAi fragments were designed, two targeting a conserved region of the *CrLFY1* and *CrLFY2* coding sequence (77% nucleotide identity) using sequences from either *CrLFY1* (*CrLFY1/2-i1*) or *CrLFY2* (*CrLFY1/2-i2*), and two targeting gene-specific sequence within the 3’UTR of *CrLFY1* (*CrLFY1-i3*) or *CrLFY2* (*CrLFY2-i4*) (**Supplementary File 6**). Target fragments were amplified from cDNA and cloned into Gateway-compatible entry vector pDONR207. Each sequence was then recombined into the pANDA expression vector via Gateway LR cloning (Invitrogen, Carlsbad, CA).

### Generation of transgenic lines

Transformation of all transgenes into wild-type Hn-n Ceratopteris callus was performed as previously described (Plackett et al., 2015). Transgenic lines were assessed in the T_1_ generation for T-DNA copy number by DNA gel blot analysis and the presence of full-length T-DNA insertions was confirmed through genotyping PCR (**Supplementary Files 5 and 6**).

### GUS staining

GUS activity analysis in *CrLFY1pro::GUS* transgenic lines was conducted in the T_1_ generation. GUS staining was conducted as described previously (Plackett et al., 2014). Optimum staining conditions (1mg/ml X-GlcA, 5uM potassium ferricyanide) were determined empirically. Tissue was cleared with sequential incubations in 70% ethanol until no further decolorization occurred.

### Phenotypic characterization

Phenotypic characterization of RNAi transgenic lines was conducted in the T_2_ or T_3_ generation. Isogenic lines were obtained by isolating hermaphrodite gametophytes in individual wells at approximately 7 DPS (or when the notch became visible, whichever came first) and flooding them once they had developed mature gametangia (at approximately 9 DPS). All transgenic lines were grown alongside wild-type controls and phenotypes observed and recorded daily. Gametophytes exhibiting altered phenotypes were imaged at approximately 10 DPS with a Nikon Microphot-FX microscope. Sporophytes with abnormal phenotypes were imaged with a dissecting microscope.

### In situ hybridization

Antisense and sense RNA probes for *CrLFY1* and *CrLFY2* were amplified and cloned into pCR 4-TOPO (Invitrogen) and DIG-labelled according to the manufacturer’s instructions (Roche, Indianapolis, IN). Probes were designed to include the 5’UTR and ORF (*CrLFY1* 521bp 5’UTR and 1113bp ORF; *CrLFY2* 301bp 5’UTR and 1185bp ORF) (**Supplementary File 7**). Tissue was fixed in FAA (3.7% formaldehyde, 5% acetic acid; 50 % ethanol) for 1-4 hours and then stored in 70% ethanol. Whole mount *in situ* hybridization was carried out based on Hejátko et al. (2006), with the following modifications: hybridization and wash steps were carried out in 24-well plates with custom-made transfer baskets (0.5 mL microcentrifuge tubes and 30 μm nylon mesh, Small Parts Inc., Logansport, IN). Permeabilization and post-fixation steps were omitted depending on tissue type, to avoid damaging fragile gametophytes, Acetic Anhydride (Sigma-Aldrich) and 0.5% Blocking Reagent (Roche) washing steps were added to decrease background staining, and tissue was hybridized at 45^°^C. Photos were taken under bright-field with a Q-imaging Micro-published 3.3 RTV camera mounted on a Nikon Microphot-FX microscope. Images were minimally processed for brightness and contrast in Photoshop (CS4).

## SUPPLEMENTAL FIGURES

**Figure 1-figure supplement 1.**
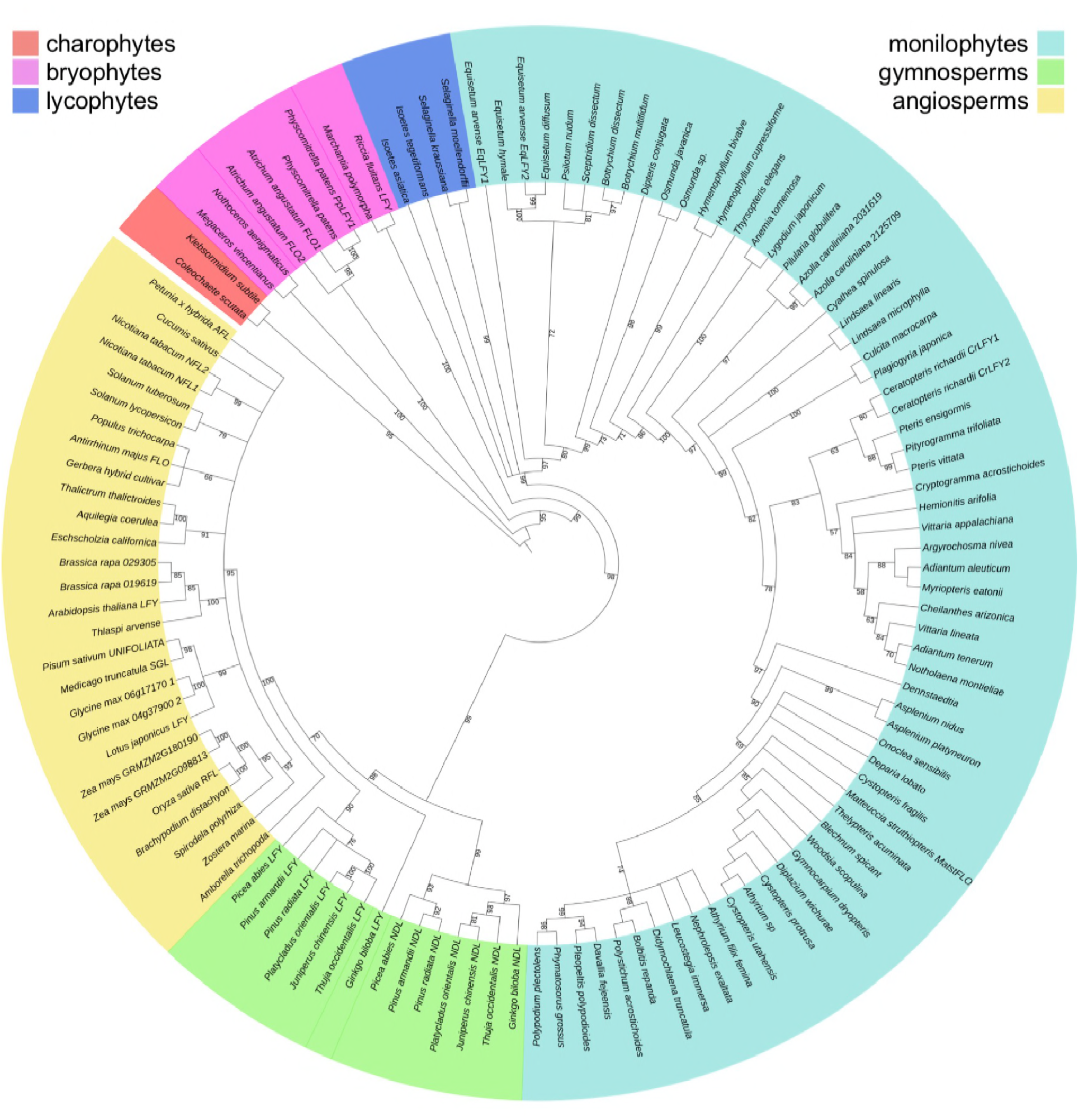
Phylogenetic relationships between *LEAFY* sequences reflect established relationships within vascular plant lineages. Inferred phylogenetic tree from maximum likelihood analysis of 120 LFY sequences sampled from across extant land plant lineages (liverworts, mosses, hornworts, lycophytes, monilophytes i.e. ferns and allies, gymnosperms, angiosperms) including algal (charophyte) sequences as an outgroup. Bootstrap values are given for each node. Sequences belonging to each lineage are denoted by different colours, as shown. The higher-order topology between vascular plant lineages (lycophytes, monilophytes, gymnosperms and angiosperms) is consistent with expected relationships; a gene duplication event resulting in *LFY* and *NEEDLY* clades in gymnosperms has been identified previously (Sayou et al., 2014); and relationships between bryophyte lineages are consistent with differences in the LFY DNA binding site preference, where hornworts and mosses each differ from the preferred site shared by liverworts and vascular plants (Sayou et al., 2014).

**Figure 1-figure supplement 2.**
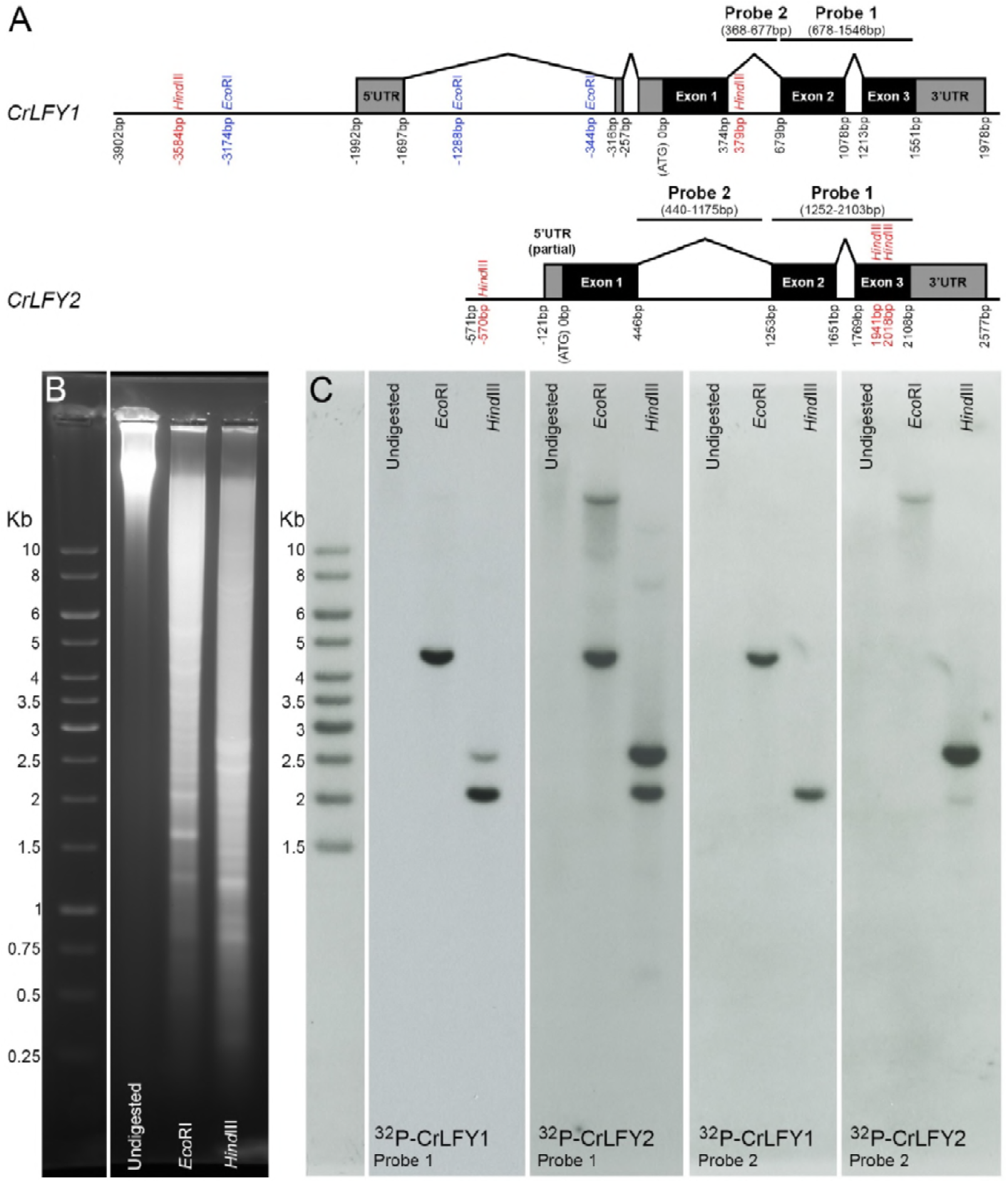
The *Ceratopteris* genome contains only two copies of *LFY*. **A**. Deduced gene structure of *CrLFY1* and *CrLFY2* loci. All positions marked are given relative to the ATG start codon. Hybridization probes used in DNA gel blot analysis and relevant restriction sites (*Eco*RI, *Hind*III) are marked. *CrLFY1* probe 1 (868bp) and *CrLFY2* probe 1 (851bp) share 79% sequence similarity and hybridize to exons 2+3 of each gene (comprising the conserved LFY DNA binding domain). As such, both probes should hybridize to all members of the *LFY* gene family. *CrLFY1* probe 2 (309bp) and *CrLFY2* probe 2 (735bp) hybridize to intron 1 of each gene copy and share no significant sequence similarity. As such, each probe is expected to hybridize to the specific gene copy. **B, C**. Gel blot analysis of wild-type genomic DNA, digested with EcoRI or HindIII, electrophoresed on an ethidium bromide stained gel (**B**), blotted to nylon membrane and hybridized against different probes (**C**) as described in (**A**). *Eco*RI digestion was predicted to generate single hybridizing fragments for both *CrLFY1* and *CrLFY2*, each spanning both probes with minimum expected fragment sizes of ~2.0kb and ~3.1kb, respectively. *Hind*III digestion was predicted to generate a single *CrLFY1* hybridizing fragment recognized by both probes with a minimum size of ~1.6kb. *Hind*III digestion was predicted to generate a *CrLFY2* fragment of ~2.5kb hybridizing to probes 1 and 2, a separate fragment with a minimum size of 559bp overlapped by 85bp of probe 1 (and so potentially undetectable) plus an undetectable fragment of 11bp. The hybridization patterns observed (**C**) are consistent with these predictions, with the exon probes cross-hybridizing to predicted fragments of both gene copies (but not to any additional gene fragments) and the intron probes primarily hybridizing to the respective specific gene copy.

**Figure 4-figure supplement 1.**
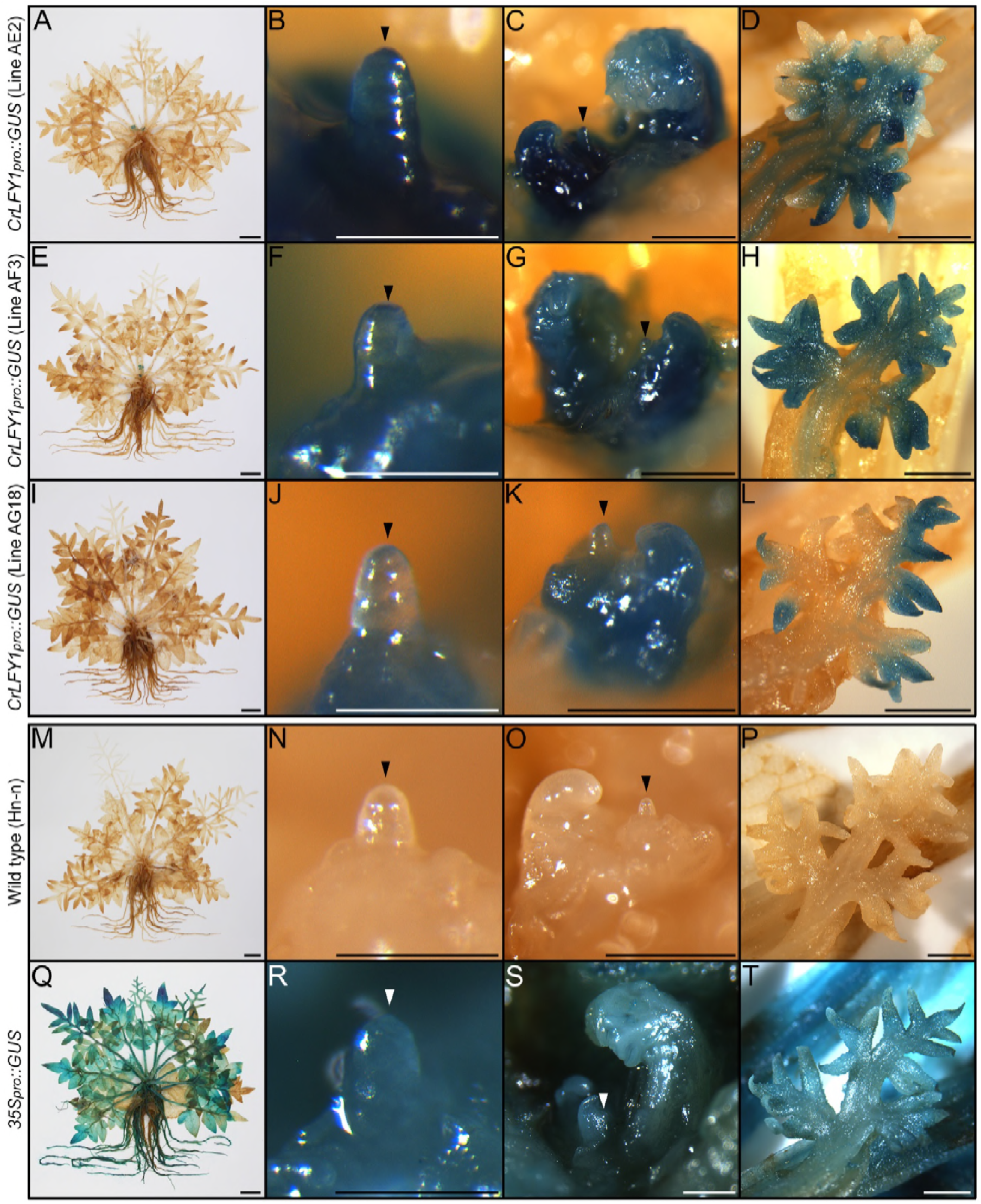
*CrLFY1_pro_::GUS* expression patterns are similar in Ceratopteris shoots before and after reproductive phase change. GUS activity detected as blue staining in sporophytes producing fronds with spore-bearing morphology (narrowing and elongation of pinnae) from three independent *CrLFY1_pro_::GUS* transgenic reporter lines (**A-L**; 110-124 DAF), negative wild-type controls (**M**-**P**; 113 DAF) and positive *35S_pro_::GUS* controls (**Q-T**; 110 DAF). Staining patterns were consistent between the three independent *CrLFY1_pro_::GUS* transgenic lines (**A, E, I**), and were similar to those seen at 60 DAF (**Figure 4 C, H, M**). GUS activity was observed throughout the shoot apex (**B, F, J**) and in recently-emerged frond primordia (**C, G, K**). Activity persisted later in frond development, becoming restricted to developing pinnae (**D, H, L**). GUS staining was lost from fronds prior to maturity (**A, E, I**). No endogenous GUS activity was detected in wild-type controls (**M-P**) whereas activity was detected throughout all non-senescent tissues in the *35S_pro_::GUS* line (**Q-T**).

**Figure 6-figure supplement 1.**
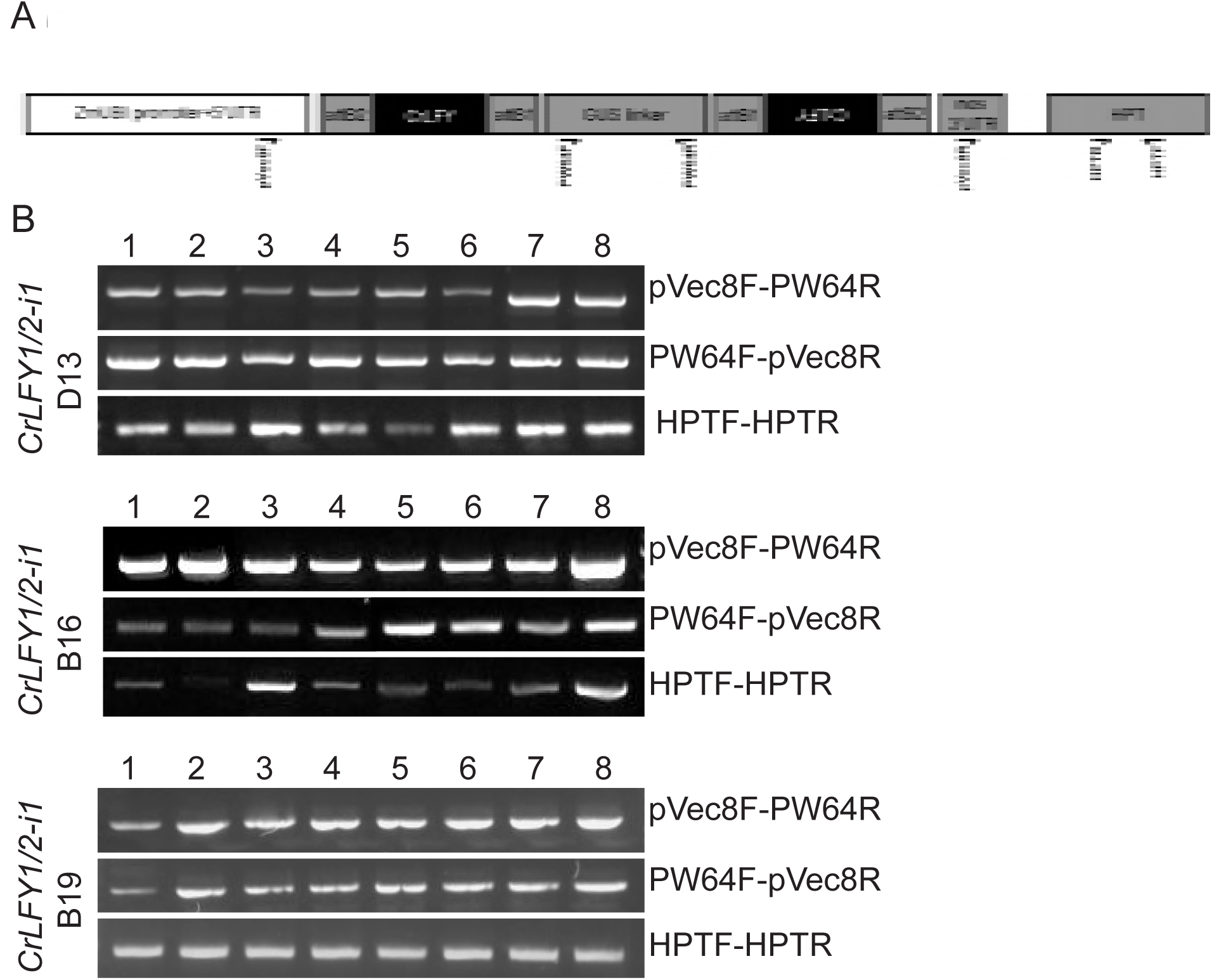
Gametophytes exhibiting developmental arrest were transgenic. **A**. Generalized schematic of *CrLFY* RNAi T-DNA with the relative position of primers used in genotyping PCR marked. Primer sequences and expected PCR product sizes for each *CrLFY* RNAi construct are given in **Table S6-8**. **B**. Genotyping PCR was conducted on DNA extracted from single gametophytes exhibiting developmental arrest at 10 DPS in three *ZmUbi_pro_::CrLFY1/2-i1* lines. DNA from all arrested individuals amplified positive bands for the two hairpin arms (pVec8F-PW64R & PW64F-pVec8R) and for the hygromycin resistance marker (HPTF-HPTR).

## SUPPLEMENTARY FILES

**Supplementary File 1.** Additional LFY sequences included in phylogenetic analysis.

**Supplementary File 2.** Alignment of all LFY amino acid sequences used in phylogenetic analysis.

**Supplementary File 3.** Alignment of LFY amino acid sequences used in phylogenetic analysis (ferns only).

**Supplementary File 4.** Statistical comparison of *CrLFY* transcript levels between different ontogenetic stages

**Supplementary File 5.** Design and validation of *CrLFY1_pro_::GUS* transgenic lines.

**Supplementary File 6.** Design and validation of *CrLFY* RNAi transgenic lines.

**Supplementary File 7.** Gel blot analysis and *in situ* hybridization probe design.

**Supplementary File 8.** Summary of published reports of LFY function in a range of angiosperm species.

AUTHOR CONTRIBUTIONS
VSD cloned the *CrLFY* coding sequences and made the RNAi constructs during a sabbatical visit to the University of Oxford; ARP and EHR performed transformations in *Ceratopteris richardii* and EHR maintained T_0_ transgenic plants; ARP cloned the *CrLFY1* promoter, made GUS reporter constructs, validated transgenic reporter lines, conducted GUS staining, and performed gel blot analysis of *CrLFY* copy number; SJC and KDHH screened, validated and characterized the RNAi lines; KDHH performed ontogenetic gene expression analysis; VSD performed statistical analyses; SJC conducted *in situ* localization experiments; JAL performed the phylogenetic analysis; JAL & ARP wrote the first draft of the paper, all authors contributed to the final version.

## ACKNOWLEDGEMENTS

Work in JAL’s lab was funded by an ERC Advanced Investigator Grant (EDIP) and by the Gatsby Charitable Foundation. VSD’s visit to JAL’s lab was funded in part by NSF/EDEN IOS # 0955517. Work in VSD’s lab was funded by the Royalty Research Fund and Bridge Funding Program, University of Washington. We thank Brittany Dean for her contribution to transgenic line screening and validation.

## REFERENCES

Ahearn KP, Johnson HA, Weigel D, Wagner DR (2001) NFL1, a Nicotiana tabacum LEAFY-like gene, controls meristem initiation and floral structure. Plant Cell Physiol 42: 1130–1139

Altschul SF, Gish W, Miller W, Myers EW, Lipman DJ (1990) Basic local alignment search tool. J Mol Biol 215: 403–410

Ambrose BA, Vasco A (2016) Bringing the multicellular fern meristem into focus. New Phytol 210: 790–793

Bar M, Ori N (2015) Compound leaf development in model plant species. Curr Opin Plant Biol 23: 61–69

Becker A, Gleissberg S, Smyth DR (2005) Floral and Vegetative Morphogenesis in California Poppy (*Eschscholzia californica* Cham.). Int J Plant Sci 166: 537–555

Bierhorst DW (1977) On the stem apex, leaf initiation and early leaf ontogeny in filicalean ferns. Am J Bot 64: 125–152

Blazquez MA, Soowal LN, Lee I, Weigel D (1997) Leafy expression and flower initiation in Arabidopsis. Development 124: 3835–3844

Bomblies K, Wang RL, Ambrose BA, Schmidt RJ, Meeley RB, Doebley J (2003) Duplicate FLORICAULA/LEAFY homologs zfl1 and zfl2 control inflorescence architecture and flower patterning in maize. Development 130: 2385–2395

Bowman JL, Sakakibara K, Furumizu C, Dierschke T (2016) Evolution in the Cycles of Life. Annu Rev Genet 50: 133–154

Bradley D, Ratcliffe O, Vincent C, Carpenter R, Coen ES (1997) Inflorescence Commitment and Architecture in Arabidopsis. Science (80-) 275: 80–83

Bradley D, Vincent C, Carpenter R, Coen E (1996) Pathways for inflorescence and floral induction in *Antirrhinum*. Development 122: 1535 LP–1544

Busch A, Gleissberg S (2003) EcFLO, a *FLORICAULA*-like gene from *Eschscholzia californica* is expressed during organogenesis at the vegetative shoot apex. Planta 217: 841–848

Carlsbecker A, Sundström Jens, Englund M, Uddenberg D, Izquierdo L, Kvarnheden A, Vergara-Silva F, Peter E (2013) Molecular control of normal and acrocona mutant seed cone development in Norway spruce (Picea abies) and the evolution of conifer ovule-bearing organs. New Phytol 200: 261–275

Carlsbecker A, Tandre K, Johanson U, Englund M, Engström P (2004) The MADS-box gene DAL1 is a potential mediator of the juvenile-to-adult transition in Norway spruce (Picea abies). Plant J 40: 546–557

Carpenter R, Coen E (1990) Floral homeotic mutations produced by transposon mutagenesis in Antirrhinum majus. Genes Dev 4: 1483–1493

Chahtane H, Vachon G, Masson M, Thévenon E, Périgon S, Mihajlovic N, Kalinina A, Michard R, Moyroud E, Monniaux M, et al (2013) A variant of LEAFY reveals its capacity to stimulate meristem development by inducing RAX1. Plant J 74: 678–689

Champagne CEM, Goliber TE, Wojciechowski MF, Mei RW, Townsley BT, Wang K, Paz MM, Geeta R, Sinha NR (2007) Compound leaf development and evolution in the legumes. Plant Cell 19: 3369–3378

Coen ES, Romero JM, Doyle S, Elliot R, Murphy G, Carpenter R (1990) floricaula: A homeotic gene required for flower development in Antirrhinum majus. Cell 63: 1311–1322

Dong Z, Zhao Z, Liu C, Luo J, Yang J, Huang W, Hu X, Wang TL, Luo D (2005) Floral Patterning in *Lotus japonicus*. Plant Physiol 137: 1272–1282

Gaillochet C, Daum G, Lohmann JU (2015) O Cell, Where Art Thou? The mechanisms of shoot meristem patterning. Curr Opin Plant Biol 23: 91–97

Ganger MT, Girouard JA, Smith HM, Bahny BA, Ewing SJ (2014) Antheridiogen and abscisic acid affect conversion and ANI1 expression in Ceratopteris richardii gametophytes. Botany 93: 109–116

Gourlay CW, Hofer JMI, Ellis THN (2000) Pea Compound Leaf Architecture Is Regulated by Interactions among the Genes UNIFOLIATA, COCHLEATA, AFILA and TENDRILLESS Plant Cell 12: 1279–1294

Harrison CJ, Morris JL (2018) The origin and early evolution of vascular plant shoots and leaves. Philos Trans R Soc B Biol Sci 373: 20160496

Hejátko J, Blilou I, Brewer PB, Friml J, Scheres B, Benková E (2006) In situ hybridization technique for mRNA detection in whole mount *Arabidopsis* samples. Nat Protoc 1: 1939–1946

Hellemans J, Mortier G, De Paepe A, Speleman F, Vandesompele J (2007) qBase relative quantification framework and software for management and automated analysis of real-time quantitative PCR data. Genome Biol 8: R19

Hickok LG, Warne TR, Fribourg RS (1995) The biology of the fern Ceratopteris and its use as a model system. Int J Plant Sci 156: 332–345

Hill JP (2001) Meristem development at the sporophyll pinna apex in Ceratopteris richardii. Int J Plant Sci 16: 235–247

Himi S, Sano R, Nishiyama T, Tanahashi T, Kato M, Ueda K, Hasebe M (2001) Evolution of MADS-box gene induction by FLO/LFY genes. J Mol Evol 53: 387–393

Hofer J, Turner L, Hellens R, Ambrose M, Matthews P, Michael A, Ellis N (1997) Unifoliata regulates leaf and flower morphogenesis in pea. Curr Biol 7: 581–588

Hou GC, Hill JP (2004) Developmental anatomy of the fifth shoot-borne root in young sporophytes of Ceratopteris richardii. Planta 219: 212–220

Johnson GP, Renzaglia KS (2008) Embryology of Ceratopteris richardii (Pteridaceae, tribe Ceratopterideae), with emphasis on placental development. J Plant Res 121: 581–592

Kanrar S, Bhattacharya M, Blake A, Courtier J, Smith HMS (2008) Regulatory networks that function to specify flower meristems require the function of homeobox genes PENNYWISE and POUND-FOOLISH in Arabidopsis. Plant J 54: 924–937

Kato M, Akiyama H (2005) Interpolation hypothesis for origin of the vegetative sporophyte of land plants. Taxon 54: 443–450

Kelly AJ, Bonnlander MB, Meeks-Wagner DR (1995) NFL, the tobacco homolog of floricaula and leafy, is transcriptionally expressed in both vegetative and floral meristems. Plant Cell 7: 225–234

Kofuji R, Hasebe M (2014) Eight types of stem cells in the life cycle of the moss Physcomitrella patens. Curr Opin Plant Biol 17: 13–21

Kyozuka J, Konishi S, Nemoto K, Izawa T, Shimamoto K (1998) Down-regulation of RFL, the FLO/LFY homolog of rice, accompanied with panicle branch initiation. Proc Natl Acad Sci U S A 95: 1979–82

Letunic I, Bork P (2016) Interactive tree of life (iTOL) v3: an online tool for the display and annotation of phylogenetic and other trees. Nucleic Acids Res 44: W242–W245

Ligrone R, Duckett JG, Renzaglia KS (2012) Major transitions in the evolution of early land plants: a bryological perspective. Ann Bot 109: 851–871

Livak KJ, Schmittgen TD (2001) Analysis of Relative Gene Expression Data Using Real-Time Quantitative PCR and the 2−ΔΔCT Method. Methods 25: 402–408

MacAlister CA, Bergmann DC (2011) Sequence and function of basic helix-loop-helix proteins required for stomatal development in Arabidopsis are deeply conserved in land plants. Evol Dev 13: 182–192

Maizel A (2005) The Floral Regulator LEAFY Evolves by Substitutions in the DNA Binding Domain. Science (80-) 308: 260–263

Maizel A, Busch MA, Tanahashi T, Perkovic J, Kato M, Hasebe M, Weigel D (2005) The floral regulator LEAFY evolves by substitutions in the DNA binding domain. Science (80-) 308: 260–263

Matasci N, Hung L-H, Yan Z, Carpenter EJ, Wickett NJ, Mirarab S, Nguyen N, Warnow T, Ayyampalayam S, Barker M, et al (2014) Data access for the 1,000 Plants (1KP) project. Gigascience 3: 17

Mellerowicz EJ, Horgan K, Walden A, Coker A, Walter C PRFLL—a Pinus radiata homologue of FLORICAULA and LEAFY is expressed in buds containing vegetative shoot and undifferentiated male cone primordia. Planta 206: 619–629

Menand B, Yi K, Jouannic S, Hoffmann L, Ryan E, Linstead P, Schaefer DG, Dolan L (2007) An ancient mechanism controls the development of cells with a rooting function in land plants. Science (80-) 316: 1477–1480

Meng Q, Zhang C, Huang F, Gai J, Yu D (2007) Molecular cloning and characterization of a *LEAFY*-like gene highly expressed in developing soybean seeds. Seed Sci Res 17: 297–302

Miki D, Shimamoto K (2004) Simple RNAi Vectors for Stable and Transient Suppression of Gene Function in Rice. Plant Cell Physiol 45: 490–495

Molinero-Rosales N, Jamilena M, Zurita S, Gómez P, Capel J, Lozano R (1999) FALSIFLORA, the tomato orthologue of FLORICAULA and LEAFY, controls flowering time and floral meristem identity. Plant J 20: 685–693

Mouradov A, Glassick T, Hamdorf B, Murphy L, Fowler B, Marla S, Teasdale RD (1998) NEEDLY, a *Pinus radiata* ortholog of *FLORICAULA/LEAFY* genes, expressed in both reproductive and vegetative meristems. Proc Natl Acad Sci 95: 6537–6542

Moyroud E, Kusters E, Monniaux M, Koes R, Parcy F (2010) LEAFY blossoms. Trends Plant Sci 15: 346–352

Moyroud E, Monniaux M, Thévenon E, Dumas R, Scutt CP, Frohlich MW, Parcy F (2017) A link between LEAFY and B-gene homologues in Welwitschia mirabilis sheds light on ancestral mechanisms prefiguring floral development. New Phytol 469–481

Nguyen L-T, Schmidt HA, von Haeseler A, Minh BQ (2015) IQ-TREE: A Fast and Effective Stochastic Algorithm for Estimating Maximum-Likelihood Phylogenies. Mol Biol Evol 32: 268–274

Niklas KJ, Kutschera U (2010) The evolution of the land plant life cycle. New Phytol 185: 27–41

Pires ND, Yi K, Breuninger H, Catarino B, Menand B, Dolan L (2013) Recruitment and remodeling of an ancient gene regulatory network during land plant evolution. Proc Natl Acad Sci. 110: 9571–9576

Plackett ARG, Huang L, Sanders HL, Langdale JA (2014) High-efficiency stable transformation of the model fern species Ceratopteris richardii via microparticle bombardment. Plant Physiol 165: 3–14

Plackett ARG, Rabbinowitsch EH, Langdale JA (2015) Protocol: Genetic transformation of the fern *Ceratopteris richardii* through microparticle bombardment. Plant Methods. 11: 37

Pnueli L, Carmel-Goran L, Hareven D, Gutfinger T, Alvarez J, Ganal M, Zamir D, Lifschitz E (1998) The self-pruning gene of tomato regulates vegetative to reproductive switching of sympodial mersitems and is the ortholog of CEN and TFL1. Development 125: 1979–1989

Proust H, Honkanen S, Jones VA, Morieri G, Prescott H, Kelly S, Ishizaki K, Kohchi T, Dolan L (2016) RSL Class I Genes Controlled the Development of Epidermal Structures in the Common Ancestor of Land Plants. Curr Biol 26: 93–99

Qiu Y-LL, Li L, Wang B, Chen Z, Knoop V, Groth-Malonek M, Dombrovska O, Lee J, Kent L, Rest J, et al (2006) The deepest divergences in land plants inferred from phylogenomic evidence. Proc Natl Acad Sci U S A 103: 15511–15516

Rao NN, Prasad K, Kumar PR, Vijayraghavan U (2008) Distinct regulatory role for RFL, the rice LFY homolog, in determining flowering time and plant architecture. Proc Natl Acad Sci 105: 3646–3651

Ratcliffe OJ, Bradley DJ, Coen ES (1999) Separation of shoot and floral identity in Arabidopsis. Development 126: 1109–1120

Rottmann WH, Meilan R, Sheppard LA, Brunner AM, Skinner JS, Ma C, Cheng S, Jouanin L, Pilate G, Strauss SH (2001) Diverse effects of overexpression of *LEAFY* and *PTLF*, a poplar (*Populus*) homolog of *LEAFY/FLORICAULA*, in transgenic poplar and *Arabidopsis*. Plant J 22: 235–245

Sakakibara K, Nishiyama T, Deguchi H, Hasebe M (2008) Class 1 KNOX genes are not involved in shoot development in the moss Physcomitrella patens but do function in sporophyte development. Evol Dev 10: 555–566

Sano R, Juárez CM, Hass B, Sakakibara K, Iyo M, Banks JA, Hasebe M, Juarez C, Hass B, Sakakibara K, et al (2005) KNOX homeobox genes potentially have similar function in both diploid unicellular and multicellular meristems, but not in haploid meristems. Evol Dev 7: 69–78

Sayou C, Monniaux M, Nanao MHH, Moyroud E, Brockington SFF, Thevenon E, Chahtane H, Warthmann N, Melkonian M, Zhang Y, et al (2014) A promiscuous intermediate underlies the evolution of LEAFY DNA binding specificity. Science (80-) 343: 645–648

Schmidt A, Schmid MW, Grossniklaus U (2015) Plant germline formation: common concepts and developmental flexibility in sexual and asexual reproduction. Development 142: 229 LP–241

Schultz EA, Haughn GW (1991) LEAFY, a Homeotic Gene That Regulates Inflorescence Development in Arabidopsis. Plant Cell 3: 771–781

Shindo S, Sakakibara K, Sano R, Ueda K, Hasebe M (2001) Characterization of a FLORICAULA/LEAFY Homologue of Gnetum parvifolium and Its Implications for the Evolution of Reproductive Organs in Seed Plants. Int J Plant Sci 162: 1199–1209

Souer E, van der Krol A, Kloos D, Spelt C, Bliek M, Mol J, Koes R (1998) Genetic control of branching pattern and floral identity during Petunia inflorescence development. Development 125: 733–742

Souer E, Rebocho AB, Bliek M, Kusters E, de Bruin RAM, Koes R (2008) Patterning of Inflorescences and Flowers by the F-Box Protein DOUBLE TOP and the LEAFY Homolog ABERRANT LEAF AND FLOWER of Petunia. Plant Cell 20: 2033–2048

Tanahashi T, Sumikawa N, Kato M, Hasebe M (2005) Diversification of gene function: homologs of the floral regulator FLO/LFY control the first zygotic cell division in the moss Physcomitrella patens Development 132: 1727–1736

Theissen G, Melzer R (2007) Molecular Mechanisms Underlying Origin and Diversification of the Angiosperm Flower. Ann Bot 100: 603–619

Theißen G, Melzer R, Rümpler F (2016) MADS-domain transcription factors and the floral quartet model of flower development: linking plant development and evolution. Development 143: 3259–3271

Tomescu AMF (2009) Megaphylls, microphylls and the evolution of leaf development. Trends Plant Sci 14: 5–12

Vasco A, Moran RC, Ambrose B (2013) The evolution, morphology, and development of fern leaves. Front Plant Sci 4: 345

Vázquez-Lobo A, Carlsbecker A, Vergara-Silva F, Alvarez-Buylla E, Piñero D, Engström P (2007) Characterization of the expression patterns of LEAFY/FLORICAULA and NEEDLY orthologs in female and male cones of the conifer genera Picea, Podocarpus, and Taxus: implications for current evo-devo hypotheses for gymnosperms. Evol Dev 9: 446–459

Wang H, Chen J, Wen J, Tadege M, Li G, Liu Y, Mysore KS, Ratet P, Chen R (2008) Control of Compound Leaf Development by FLORICAULA/LEAFY Ortholog SINGLE LEAFLET1 in Medicago truncatula. Plant Physiol 146: 1759–1772

Weigel D, Alvarez J, Smyth DR, Yanofsky MF, Meyerowitz EM (1992) LEAFY controls floral meristem identity in Arabidopsis. Cell 69: 843–859

Weigel D, Nilsson O (1995) A developmental switch sufficient for flower initiation in diverse plants. Nature 377: 495–500

White R, Turner M (1995) Anatomy and development of the fern sporophyte. Bot Rev 61: 281–305

Wickett NJ, Mirarab S, Nguyen N, Warnow T, Carpenter E, Matasci N, Ayyampalayam S, Barker MS, Burleigh JG, Gitzendanner MA, et al (2014) Phylotranscriptomic analysis of the origin and early diversification of land plants. Proc Natl Acad Sci 111: E4859–E4868

Wreath S, Bartholmes C, Hidalgo O, Scholz A, Gleissberg S (2013) Silencing of EcFLO, A FLORICAULA/LEAFY Gene of the California Poppy (*Eschscholzia californica*), Affects Flower Specification in a Perigynous Flower Context. Int J Plant Sci 174: 139–153

Yang T, Du MF, Guo YH, Liu X (2017) Two LEAFY homologs ILFY1 and ILFY2 control reproductive and vegetative developments in Isoetes L. Sci Rep. 1: 225

Yip HK, Floyd SK, Sakakibara K, Bowman JL (2016) Class III HD-Zip activity coordinates leaf development in Physcomitrella patens. Dev Biol 419: 184–197

Zhao W, Chen Z, Liu X, Che G, Gu R, Zhao J, Wang Z, Hou Y, Zhang X (2017) CsLFY is required for shoot meristem maintenance via interaction with WUSCHEL in cucumber (Cucumis sativus). New Phytol 218: 344–356

Zhao Y, Zhang T, Broholm SK, Tähtiharju S, Mouhu K, Albert VA, Teeri TH, Elomaa P (2016) Evolutionary Co-Option of Floral Meristem Identity Genes for Patterning of the Flower-Like Asteraceae Inflorescence. Plant Physiol 172: 284–296

